# mfSuSiE enables multi-cell-type fine-mapping and multi-omic integration of chromatin accessibility QTLs in aging brain

**DOI:** 10.1101/2025.11.25.690439

**Authors:** Anjing Liu, Hao Sun, Philip L. De Jager, David Bennett, The Alzheimer’s Disease Functional Genomics Consortium, Gao Wang, William R. P. Denault

## Abstract

Molecular quantitative trait locus (QTL) studies increasingly profile chromatin accessibility, histone modifications, DNA methylation, RNA modifications such as N6-methyladenosine (m6A), and transcription across multiple cell types using high-throughput sequencing, generating dense base-pair-resolved measurements. The conventional approach of testing each variant against each molecular feature independently suffers from severe multiple testing burden and ignores linkage disequilibrium and spatial correlation. Existing fine-mapping methods only partially address these challenges and are sub-optimal for analyzing such datasets: multivariate approaches such as *mvSuSiE* jointly analyze multiple molecular contexts but are designed for a single trait value per context and cannot accommodate thousands of base-resolution measurements per context, while functional approaches such as *fSuSiE* model spatial structure across thousands of measurements but analyze each context separately. Here, we introduce *mfSuSiE*, which integrates multivariate analysis with wavelet-based functional regression to jointly fine-map thousands of base-resolution traits across multiple cell types. In simulations, *mfSuSiE* identified causal variants and affected molecular features more accurately than *fSuSiE*, while *mvSuSiE* cannot be applied to this type of data. Applied to single-nucleus chromatin accessibility data from six brain cell types from postmortem aging human brains, *mfSuSiE* substantially increased discovery and resolution, with substantial power gains for cell types with limited samples. Multi-cell-type analysis revealed extensive sharing of regulatory effects on chromatin accessibility (caQTL). Importantly, *mfSuSiE* produces Bayesian inference compatible with the *SuSiE* framework, enabling systematic multi-omic integration. Applied to Alzheimer’s disease loci, we integrated caQTL with expression QTLs, epigenomic QTLs, and GWAS, observing regulatory patterns suggesting complex mechanisms at loci including *EARS2, CHRNE, SCIMP*, and *RABEP1*.

## INTRODUCTION

The analysis of molecular quantitative trait loci (QTLs) has advanced rapidly over the past decade. Early studies typically examined associations between individual genetic variants and single molecular traits measured in one tissue or cell type. However, large-scale multi-tissue projects have shown that many genetic effects are shared across related tissues and molecular modalities [1, 2]. These findings have spurred the development of statistical methods that jointly analyze multiple correlated traits. Existing fine-mapping approaches, such as *SuSiE* [3] and CAVIAR [4] for univariate traits, or *mvSuSiE* [2] and CAFEH [5] for multivariate traits, have improved the resolution at which causal variants can be identified. Yet these models treat traits as vectors of independent measurements — for example, expression levels across tissues — without explicitly modeling the rich spatial structure captured by modern sequencing-based molecular assays.

High-throughput assays such as ATAC-seq, RNA-seq, and ChIP-seq provide dense, base-pair–resolved profiles along the genome, revealing local regulatory architectures and spatial patterns in chromatin accessibility, transcription, and protein binding. We previously introduced functional *SuSiE* (*fSuSiE*) [6, 7] to show that these high-resolution molecular readouts are more naturally represented as functions over genomic coordinates than as isolated, position-wise summary statistics. Functional representations better capture the biological reality that causal genetic variants often induce spatially structured changes, such as methylation QTLs affecting clusters of CpG sites, or splicing QTLs altering read-depth profiles across introns.

Despite this progress, a critical gap remains in understanding how genetic variants shape molecular phenotypes across multiple tissues or cell types simultaneously. Growing multi-omic datasets now provide matched sequencing-based molecular profiles for several cell types within the same individuals, creating an opportunity to dissect shared versus context-specific regulatory mechanisms. The ability to jointly analyze chromatin accessibility across multiple cell types and integrate these results with corresponding gene expression profiles would enable more precise characterization of transcriptional regulatory mechanisms underlying complex traits. Because many genetic effects on complex traits are mediated by cell-type-specific regulatory variation, and because regulatory effects often manifest across multiple molecular layers (chromatin accessibility, gene expression, histone modifications), the absence of methods that can jointly model multivariate and spatially structured molecular traits while facilitating integration across modalities has become a major barrier to interpreting disease-associated loci.

Here, we introduce multivariate functional *SuSiE* (*mfSuSiE*), a fine-mapping framework that extends *fSuSiE* to jointly model multiple functional molecular phenotypes across tissues, cell types, or modalities. *mfSuSiE* represents the effect of each candidate causal variant as a multivariate function along the genome, enabling the discovery of effects that are shared, partially shared, or cell-type-specific. Building on principles from functional data analysis [6, 8–21], *mfSuSiE* retains the computational scalability of *SuSiE* [3, 22] by decomposing multivariate fine-mapping into a sequence of tractable single-effect updates. This design allows *mfSuSiE* to scale to tens of thousands of genomic positions, dozens of tissues, and thousands of candidate variants. A key advantage of embedding our approach within the *SuSiE* framework is seamless compatibility with the growing ecosystem of *SuSiE*- based analytical tools, enabling sophisticated multi-omic integration pipelines through colocalization analysis and other downstream methods.

We evaluate *mfSuSiE* using simulated multi-context functional QTL data and show that joint modeling substantially improves fine-mapping accuracy compared to applying *fSuSiE* or univariate methods independently to each context. We then apply *mfSuSiE* to chromatin accessibility profiles from six major human brain cell types (astrocytes, excitatory neurons, inhibitory neurons, microglia, oligodendrocytes, and oligodendrocyte precursor cells) generated by single-nucleus ATAC-seq from the Religious Orders Study (ROS) and Memory and Aging Project (MAP), a longitudinal cohort study of aging and Alzheimer’s disease (AD) [23]. Through follow-up colocalization analysis, we integrate chromatin accessibility (caQTL) results with expression QTLs (eQTLs) on single-nucleus RNA-seq from the same cell types, as well as bulk brain eQTL and epigenomic QTLs (histone modification and DNA methylation), and AD GWAS fine-mapping. We demonstrate how this multi-omic integration refines variant- and gene-prioritization and elucidates regulatory mechanisms at AD risk loci.

## RESULTS

### Overview of Method

*mfSuSiE* extends *fSuSiE* [24] to jointly fine-map multiple molecular modalities. It takes as input a collection of matrices (**Y**_*m*_)_*m*=1,…,*M*_, where each **Y**_*m*_ is an *N* × *T*_*m*_ matrix containing a molecular phenotype for *N* samples measured at *T*_*m*_ genomic locations. Let **X** denote the *N* × *P* genotype matrix genotypes for the same samples at *P* candidate genetic variants. *mfSuSiE* models each modality by a multivariate functional linear model,

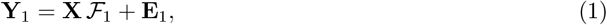

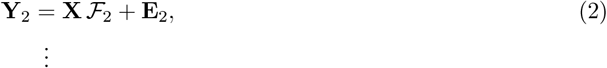

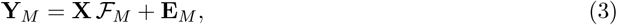

where ℱ_*m*_ is a *P* × *T*_*m*_ matrix of spatially varying effects for modality *m*, and **E**_*m*_ is a matrix of residuals. The element ℱ_*m,jt*_ represents the effect of SNP *j* on genomic position *t* in modality *m*.

As in most fine-mapping methods [3, 25], *mfSuSiE* assumes that only a small number of SNPs have non-zero effects, so most rows of ℱ_*m*_ are zero. To impose this sparsity, *mfSuSiE* uses a multivariate “sum of single effects” (*SuSiE*) decomposition [3, 7, 22]:

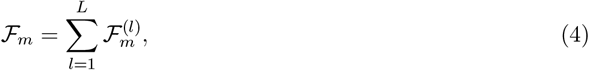

where each component 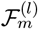 corresponds to the effect of a single causal variant and therefore contains exactly one non-zero row. Crucially, the non-zero row is shared across all modalities for a given *l*, ensuring that each component identifies a s hared c ausal S NP w hile a llowing i ts f unctional e ffect to differ across modalities.

To model spatial structure efficiently, *mf SuSiE* adopts the discrete wavelet representation used in *fSuSiE* [12, 17, 26, 27]. Wavelets provide a compact multiscale representation of molecular phenotypes, capturing both sharp local changes and broad genomic domains. In practice, *mfSuSiE* first applies a wavelet transform to each modality in (1), then applies sparse priors to the transformed effect coefficients to impose shrinkage.

A key practical advantage of *mfSuSiE* is that it naturally handles missing data. Individuals may lack measurements for certain modalities—for example, a donor may have ATAC-seq profiled in astrocytes and oligodendrocytes but not in excitatory neurons. *mfSuSiE* performs inference using all available observations without requiring imputation or restricting to complete cases.

Further, *mfSuSiE* accommodates mixtures of univariate and functional phenotypes, such as scalar expression values for one modality and spatially resolved ATAC-seq profiles f or a nother. For clarity, we present the case where all modalities share the same genomic grid (*T*_1_ = *…* = *T*_*M*_ = *T*), although the framework can in principle support modalities defined over distinct genomic regions.

### Numerical study

We evaluated the performance of *mfSuSiE* through a series of simulation studies designed to reflect effect of SNPs on one or more molecular trait (Figure 2. Genotypes were simulated using `sim1000G` [28] with 1000 Genomes Phase 3 whole-genome sequencing data as input, yielding genomic regions of approximately 1 Mb containing 1,500–4,000 SNPs. These regions exhibited realistic patterns of linkage disequilibrium (LD), including cases of strong or perfect correlation between SNPs, making exact causal SNP identification inherently challenging. Functional molecular phenotypes were simulated following the wavelet-based generative model introduced in our previous work (*fSuSiE* “wavelet simulations”), while univariate phenotypes were generated according to the model of [3]. Across simulation replicates, we varied the number of SNPs, the number of causal variants, and the noise level. Detailed descriptions of the simulation settings are provided in the Online Methods.

**Fig. 1.**
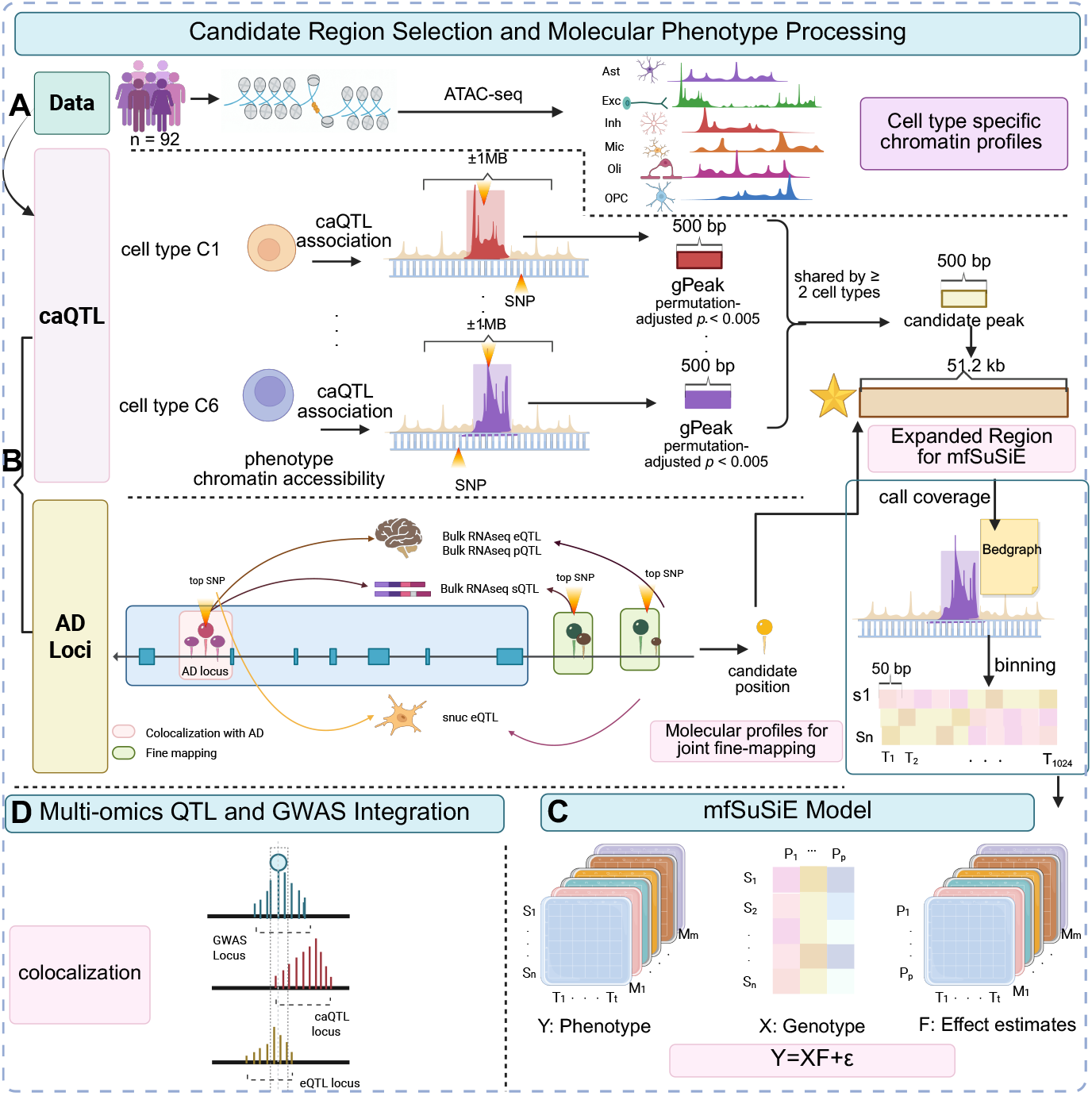
Overview of *mfSuSiE* and multi-cell-type chromatin accessibility QTL fine-mapping. Each panel illustrates a component of the fine-mapping framework for multi-cell-type single-nucleus ATAC-seq data: **(A)** Data processing from ROSMAP snATAC-seq (n=92 samples) generating cell-type-specific chromatin accessibility profiles across six brain cell types (Ast: Astrocytes, Exc: Excitatory neurons, Inh: Inhibitory neurons, Mic: Microglia, Oli: Oligodendrocytes, OPC: Oligodendrocyte precursor cells). **(B)** Candidate region selection: 156 established AD GWAS risk loci identified through FunGen-xQTL multi-context colocalization, plus 12 regions where marginal caQTL associations (gPeaks, permutation-adjusted *p* < 0.005) were shared across at least two cell types. Each region is expanded to 51.2kb windows. Phenotype processing: base-level ATAC-seq coverage is binned into 1,024 bins × 50bp (51.2kb per region), creating high-resolution chromatin accessibility matrices for *mfSuSiE* fine-mapping. **(C)** *mfSuSiE* joint modeling across six cell types, enabling simultaneous fine-mapping to identify shared and cell-type-specific causal variants. **(D)** Multi-omic integration enabled by *mfSuSiE*, including colocalization with expression QTLs, epigenomic QTLs, and AD GWAS to characterize regulatory mechanisms at disease loci.

**Fig. 2.**
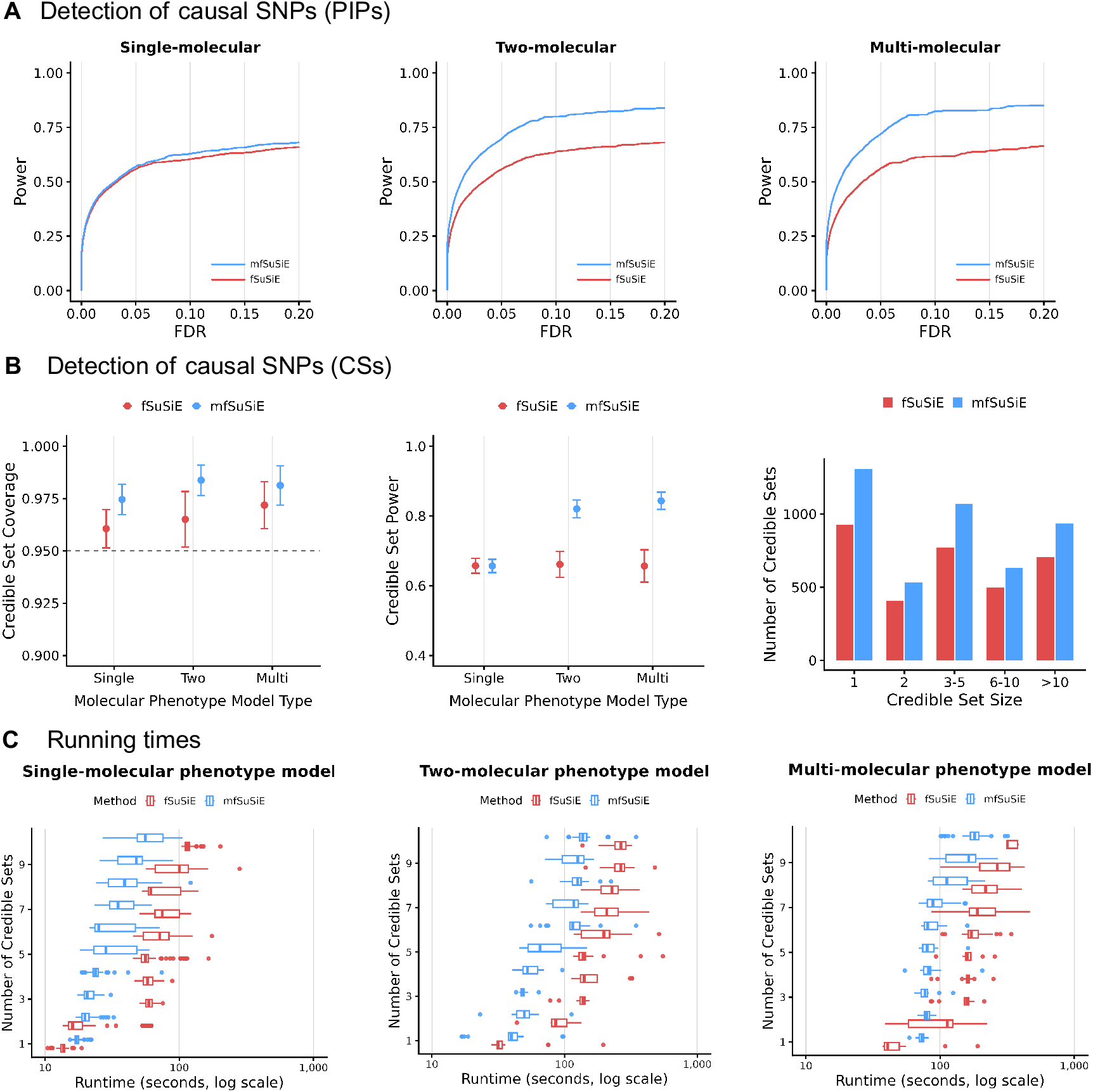
*mfSuSiE* simulation study: power, coverage, and computational efficiency. Simulation evaluation of method performance across three modeling scenarios: single-molecular phenotype model, two-molecular phenotype model, and multi-phenotype model. All simulations used sample size *N* = 100, with each simulation randomly selecting 1–20 causal variants. The single-molecular phenotype model contains single molecular phenotype (32 columns). The two-molecular phenotype model includes 2 molecular phenotypes (32 + 64 columns), showing improved power through joint modeling. The multi-phenotype model includes 2 molecular phenotypes + 3 univariate phenotypes, demonstrating benefits of integrating diverse data modalities. **(A) Detection of causal SNPs (PIPs)**. FDR and power curves for the three modeling scenarios under high noise (SD=3). Blue and red curves represent *mfSuSiE* and *fSuSiE* methods, respectively. **(B) Detection of causal SNPs (CSs)**. Evaluation under high noise (SD=3). Left panel shows credible set coverage, assessing whether credible sets contain true causal variants at nominal coverage. Middle panel shows credible set power across model complexities. Right panel displays the number of credible sets by size for the two-molecular phenotype model. **(C) Running times**. Computational efficiency comparison across methods for the three modeling scenarios, showing runtime across varying noise levels (SD=1 to SD=4).

To compare *mfSuSiE* and *fSuSiE*, we considered three modeling scenarios under a fixed sample size of *N* = 100: a single functional phenotype analyzed by both methods; two molecular phenotypes analyzed jointly by *mfSuSiE* versus fine mapping separately both molecular by *fSuSiE* and reports CS for both; and a full model in which *mfSuSiE* jointly analyzes two functional and three univariate phenotypes, while *fSuSiE* analyzes only the first functional phenotype. This design allowed us to assess the benefits of multivariate joint modeling relative to single-trait functional fine-mapping.

Across all scenarios, we examined the methods’ ability to identify causal SNPs affecting at least one molecular trait at one or more genomic locations. Both methods produce posterior inclusion probabilities (PIPs) and 95% credible sets (CSs), which serve as the primary outputs in fine-mapping analyses. Because LD often makes it impossible to pinpoint a single causal SNP, Credible sets provide a more appropriate metric of performance: a method is considered effective if it returns small, high-purity CSs with reliable coverage (i.e., the proportion of true SNP in a given CS, target coverage = 95%). While PIPs remain informative for ranking candidate variants, CS size, coverage, and purity are the most relevant measures of fine-mapping accuracy.

Unsurprisingly, *mfSuSiE* and *fSuSiE* performed similarly when applied to a single functional phenotype. However, the benefits of *mfSuSiE* became apparent once multiple phenotypes were introduced. In the two-phenotype and full-model settings, *fSuSiE* (single-phenotype analysis) exhibited reduced sensitivity, and diminished power. In contrast, *mfSuSiE* effectively leveraged information across modalities, producing smaller and more precise credible sets while maintaining high coverage and distinguishing true causal variants more reliably. These results highlight the advantages of joint functional fine-mapping when causal variants exert shared or partially shared effects across molecular traits.

### Application to chromatin accessibility QTLs in aging brain

To demonstrate the utility of multi-cell-type functional fine-mapping, we applied *mfSuSiE* to single-nucleus ATAC-seq data from the ROSMAP study [23]. Chromatin accessibility profiles were obtained from six major brain cell types previously profiled in 92 donors from dorsolateral prefrontal cortex [29], including astrocytes (Ast), excitatory neurons (Exc), inhibitory neurons (Inh), microglia (Mic), oligodendrocytes (Oli), and oligodendrocyte precursor cells (OPC).

We selected 156 established Alzheimer’s disease GWAS risk loci identified through multi-context xQTL colocalization by the FunGen-xQTL project, with the goal of characterizing cell-type-specific chromatin accessibility regulation at disease loci of interest. We additionally included 12 regions where conventional marginal association testing detected shared signals across at least two cell types, to assess whether *mfSuSiE* could recover effects detectable by standard approaches. We detail the selection of these regions in online Methods section *Molecular QTL data resource and processing*. For each region, we constructed high-resolution chromatin accessibility phenotypes by binning base-level coverage into 1,024 genomic positions spanning 51.2 kb windows (50 bp bins), and extracted genotype data for SNPs within *±*500 kb (typically 2,000–4,000 SNPs per region) from whole-genome sequences on these donors [30]. Data processing and quality control procedures are detailed in the **Methods**.

Multi-cell-type fine-mapping with *mfSuSiE* identified 609 candidate regulatory variants from a total of 332 CS (target coverage (5%) across candidate regions after quality control filtering, following standard fine-mapping practices established in the FunGen-xQTL project (variants within CS having purity *>* 0.8, or outside of CS with posterior inclusion probability *>* 0.1). Compared to single-cell-type analysis with *fSuSiE*, joint modeling with *mfSuSiE* across cell types produced more credible sets overall and notably more compact credible sets, achieving higher resolution in prioritizing candidate causal variants (Fig. 3A). To characterize which genomic positions and cell types are affected by each credible set, we computed local false sign rates (LFSR) for the functional regression coefficients at each genomic position and cell type; positions with LFSR < 0.01 were considered to show significant effects. Among credible sets meeting quality thresholds, *mfSuSiE* identified substantially more with significant effects across cell types compared to *fSuSiE* (Fig. 3B). The improvement was particularly significant for oligodendrocyte precursor cells: single-cell-type *fSuSiE* identified essentially no credible sets with significant OPC effects, whereas *mfSuSiE* identified over 200 such credible sets by leveraging shared regulatory architecture with other brain cell types. Beyond providing smaller credible sets and increased discovery, *mfSuSiE* maintains strong consistency with single-cell-type *fSuSiE* results (Fig. 3C): the majority of *fSuSiE* credible sets (90.2%) showed direct correspondence with an *mfSuSiE* credible set, demonstrating that *mfSuSiE* robustly replicates signals identified by single-cell- type analysis.

**Fig. 3.**
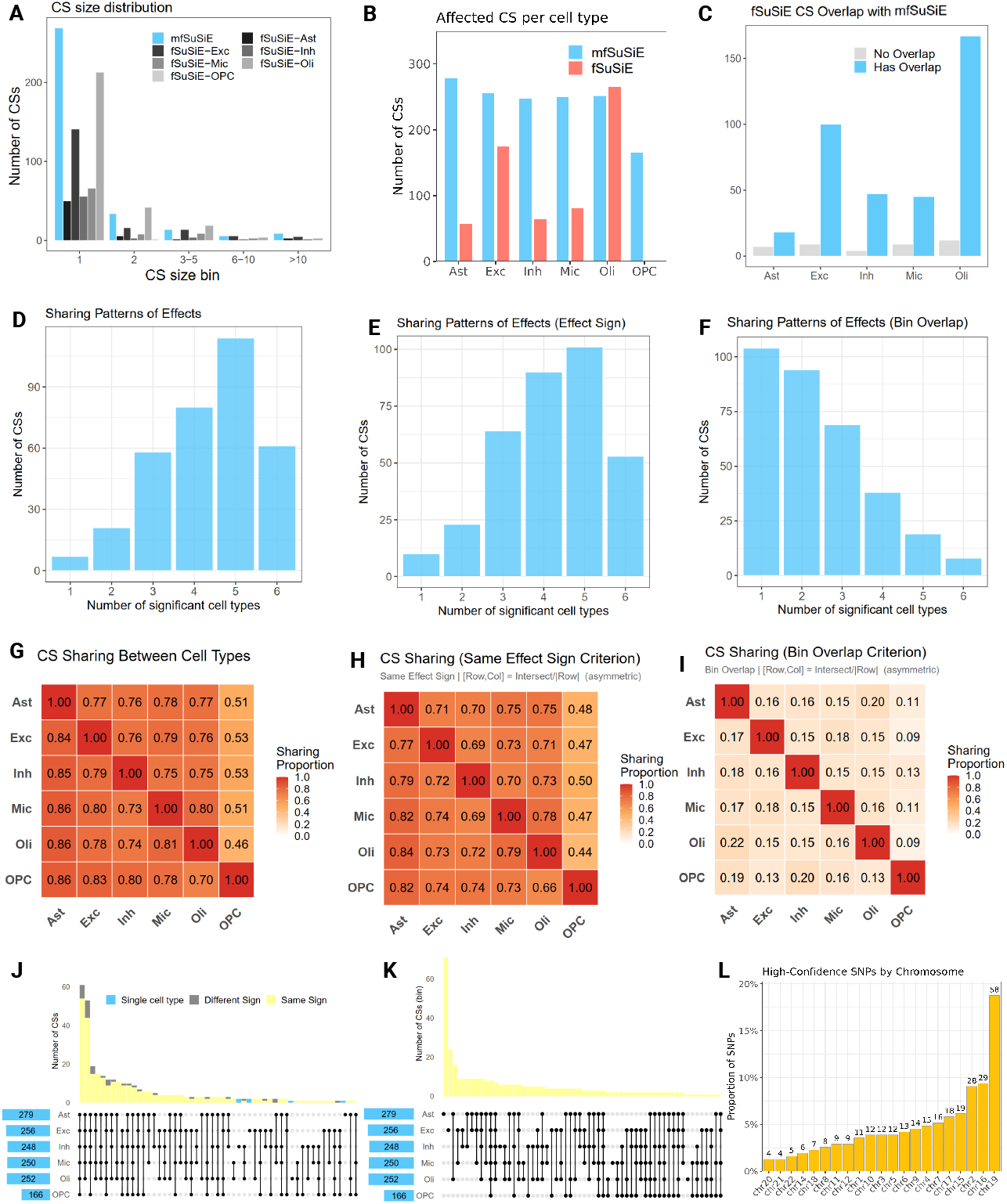
Overview of *mfSuSiE* fine-mapping results and comparison with single-cell-type analysis. All panels summarize results from *mfSuSiE* or single-cell-type *fSuSiE* under consistent filtering criteria. Credible sets (CSs) are retained when purity exceeds 0.8, where purity is the minimum absolute correlation among variants within a CS. Posterior inclusion probability (PIP) reflects the posterior probability that each variant is causal; variants with PIP *>* 0.1 outside 95% CSs are included as additional candidates. Local false sign rate (LFSR) is the posterior probability of incorrect effect direction; LFSR < 0.01 indicates significant effects. Bin-overlap is defined as sharing a significant effect (LFSR < 0.01) in the same 50 bp genomic bin within a *±*200 bp window across cell types. **(A)** Distributions of CS sizes for *mfSuSiE* (blue) and single-cell-type *fSuSiE* (gray) after purity filtering. **(B)** Number of CSs per cell type containing at least one SNP with LFSR < 0.01. **(C)** Overlap of *fSuSiE* CSs with *mfSuSiE* CSs, stratified into overlapping and non-overlapping sets. **(D–F)** Sharing patterns of CSs across six brain cell types under three criteria. (D) Basic sharing criterion: CSs with purity *>* 0.8 and lowest LFSR < 0.01 in each cell type; bars show the number of CSs that are significant in exactly *k* cell types. (E) Same-effect-sign criterion: restricted to CSs passing the basic criterion, further requiring that more than 50% of effects across cell types have the same sign. (F) Bin-overlap criterion: CSs with at least one bin (LFSR < 0.01) shared across multiple cell types. **(G–I)** Asymmetric pairwise CS sharing between cell types under the same three criteria. Entries show, for each row cell type, the fraction of its CSs that are shared with the column cell type; sharing is defined by the corresponding basic, same-effect-sign, or bin-overlap rule, and computed as the size of the intersection divided by the number of CSs in the row cell type. **(J–K)** UpSet-style summaries of multi-cell-type CS sharing. (J) Combined view under basic and same-effect-sign criteria. (K) Sharing under the bin-overlap criterion. Bars display the number of CSs shared by each combination of cell types. **(L)** Chromosome distribution of putative causal variants (PIP *>* 0.5) within CSs meeting the purity threshold.

Analysis of these credible sets with significant effects revealed extensive sharing of regulatory architecture. The distribution of significant cell-type effects peaked at 5–6 cell types per credible set (Fig. 3D), indicating that most regulatory variants affect chromatin accessibility across multiple brain cell lineages rather than acting in a single cell type. Applying additional constraints on effect direction and spatial concordance progressively reduced the extent of sharing but preserved the overall pattern (Fig. 3E–F). Directional consistency across cell types moderately attenuated sharing, while requiring spatial overlap of functional effects yielded a more conservative subset of shared signals, reflecting tighter regulatory coordination. Pairwise comparisons of credible set sharing confirmed high concordance among major neuronal (Exc, Inh) and glial (Ast, Mic) populations, while OPC exhibited more distinctive patterns with other cell types; Fig. 3G-I), consistent with its specialized developmental role. UpSet representations provided a complementary summary of these results. The UpSet plot based on significance and directional consistency (Fig. 3J) confirmed that the most frequent sharing configurations involved simultaneous effects across multiple neuronal (Exc, Inh) and glial (Ast, Mic) cell types, consistent with a broadly shared regulatory architecture. In contrast, the spatial overlap–based UpSet plot (Fig. 3K) exhibited markedly sparser configurations, dominated by lower-order overlaps, indicating that precise spatial alignment of regulatory effects is restricted to a smaller subset of cell-type combinations. Genome-wide distribution of putative causal variants (PIP *>* 0.5 within credible sets) showed that chromosome 19 had the highest concentration (18.5%, n=58 variants), mapping predominantly to the *APOE* /*TOMM40* /*APOC* cluster, and chromosome 16 also showed notable enrichment (9.5%, n=29 variants), largely within the *DOC2A*/*YPEL3* /*INO80E* cluster (Fig. 3L). These two regions are among the most gene-dense among AD risk loci, so the concentration of regulatory variants in these regions is consistent with their dense regulatory architecture. However, both regions also exhibit extensive linkage disequilibrium, which may complicate interpretation of independent causal signals.

### Multi-omic integration via colocalization analysis

One added advantage of fine-mapping chromatin accessibility QTLs with *mfSuSiE* is the ability to integrate these results with other molecular QTL data to strengthen discoveries and provide mechanistic insights, addressing the inherent challenge of limited sample sizes in some molecular QTL studies (in this case, our single-nucleus ATAC-seq data). Recent advances in colocalization analysis enable leveraging fine-mapping results directly as input [31]; extending this framework allows us to use *mfSuSiE* credible sets to test for shared genetic effects with expression QTLs and GWAS, combining evidence across complementary data types.

We performed colocalization analysis to integrate our *mfSuSiE* chromatin accessibility results with additional FunGen-xQTL data. Expression QTLs were fine-mapped using *mvSuSiE* [32] on bulk and single-nucleus RNA-seq from the same ROSMAP donors (n=416 bulk; cell-type sample sizes in **Methods**), analyzing the six matched brain cell types plus additional bulk contexts. Each method, *mvSuSiE* (multivariate fine-mapping with adaptive shrinkage mixture prior) and *mfSuSiE* (wavelet-based functional fine-mapping across multi-traits), was designed for its specific molecular context to characterize cell-type sharing patterns, boost statistical power, and improve detection accuracy. Combining them enables 12-dimensional integration: six cell types with paired chromatin accessibility and gene expression, each fine-mapped in its appropriate framework. We further integrated with Alzheimer’s disease GWAS fine-mapping (using *SuSiE* on summary statistics [33]) to focus our analysis on established disease loci rather than genome-wide discovery. This multi-modal, multi-cell-type integration framework has not been previously demonstrated and is made possible by SuSiE-based fine-mapping compatibility. Applying this framework, we illustrated candidate regulatory landscape at two loci, described below.

#### Multi-omic convergence at a suggestive AD risk region with *EARS2*

Colocalization analysis between chromatin accessibility fine-mapping and AD GWAS (*mfSuSiE*–*SuSiE*) identified a suggestive association at the *EARS2* locus (chr16:23.4–23.5 Mb, peak GWAS *p* = 2.41 × 10^−6^), yielding a credible set of 33 variants (PP.H4 *>* 0.8). We also performed colocalization between chromatin accessibility and expression QTLs (*mfSuSiE*–*mvSuSiE*), identifying a credible set of 19 variants. The overlap between these two colocalization analyses prioritized a one-base deletion at chr16:23580183, showing significant effects on chromatin accessibility in oligodendrocytes, astrocytes, microglia, and excitatory neurons (LFSR < 0.01) and on *EARS2* expression in excitatory neurons, inhibitory neurons, oligodendrocytes (cell-type-specific eQTLs, LFSR < 0.01), plus dorsolateral prefrontal cortex, posterior cingulate cortex, and anterior cingulate cortex (bulk eQTLs, LFSR < 0.01).

Integration with expression QTL data revealed consistent regulatory effects across molecular layers. The chromatin accessibility increases associated with the candidate variant corresponded to elevated *EARS2* expression in both bulk tissue and cell-type-specific contexts. Functional annotation of the 19 variants in *mvSuSiE*–*mfSuSiE* colocalization demonstrated enrichment for active regulatory features: 12 of examined variants fell within a super-enhancer region marked by H3K27ac, 9 showed H3K4me1 enhancer marks, and several exhibited evidence of transcription factor binding and RNA polymerase II occupancy. Hi-C interaction data indicated long-range enhancer-promoter contacts, supporting direct regulatory communication between the variant-containing region and the *EARS2* promoter. Complete functional annotations are provided in Supplementary Table 3.

*EARS2* encodes glutamyl-tRNA synthetase 2, essential for mitochondrial protein synthesis. Disruption of *EARS2* function leads to mitochondrial dysfunction, which has been implicated in neurodegenerative processes. While *EARS2* has not been definitively linked to AD pathogenesis, the convergent evidence from chromatin accessibility, gene expression, and AD GWAS, combined with its biological role in mitochondrial function known to regulate amyloid-beta and tau proteins in AD, suggests this locus warrants further investigation as a potential contributor to AD risk through mitochondrial mechanisms.

#### Coordinated regulation across genomic distance at AD risk loci involving multiple genes

Colocalization analysis at an established AD risk locus on chromosome 17 (chr17:4.8–5.3 Mb, peak GWAS *p* = 6.84 × 10^−13^) revealed complex regulatory architecture involving multiple genes. This region contains a cluster including *CHRNE* (cholinergic receptor nicotinic epsilon subunit), *SCIMP* (SLP adaptor and CSK interacting membrane protein), and *RABEP1* (rabaptin, RAB GTPase binding effector protein 1). Colocalization between chromatin accessibility fine-mapping and AD GWAS (*mfSuSiE*–*SuSiE*) identified variants at approximately 4.9 Mb (PP.H4 = 0.86).

Examining the fine-mapped chromatin accessibility effects more closely, we found that variants at the 4.9 Mb position affect chromatin accessibility across all six brain cell types, with significant effects (LFSR < 0.01) extending to the 5.2 Mb region (Fig. 6). These distal effects span approximately 330 kb and encompass the *SCIMP* gene region, including 3.8 kb upstream and 21.4 kb downstream of the gene body, but do not extend to *RABEP1* (located 43.6 kb away). Fine-mapping with *mvSuSiE* identified expression QTL credible sets at the 5.2 Mb region for both *SCIMP* (microglia, astrocytes, excitatory neurons, inhibitory neurons, and bulk tissues) and *RABEP1* (microglia, oligodendrocytes, and bulk tissues). The target regions of chromatin accessibility effects from the 4.9 Mb variants overlap with these fine-mapped eQTL credible sets in their respective cell types, confirming functional consequences of the distal chromatin changes (Fig. 6).

**Fig. 4.**
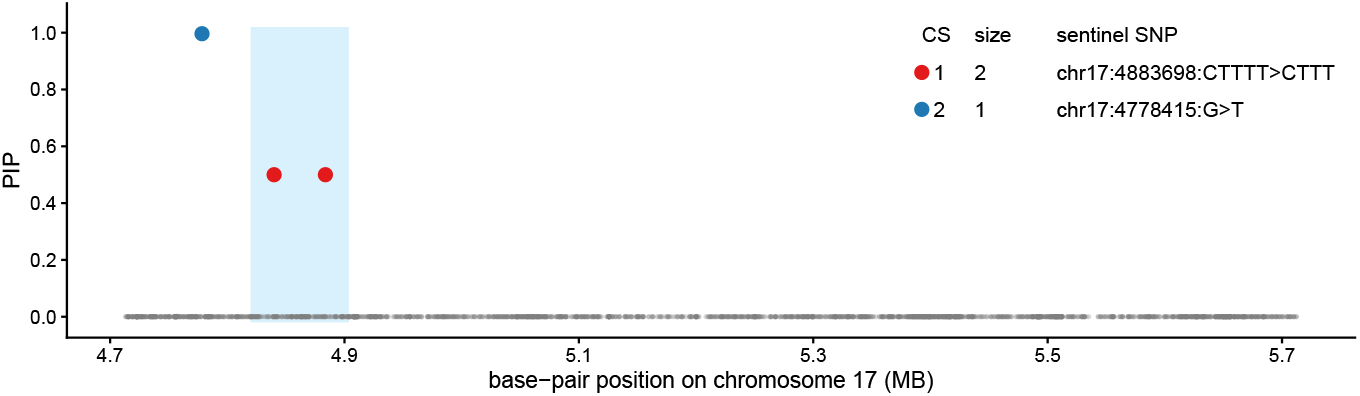
Example *mfSuSiE* fine-mapping of chromatin accessibility. Posterior inclusion probabilities (PIPs) for all SNPs analyzed in the region chr17:4,713,084–5,713,084.

**Fig. 5.**
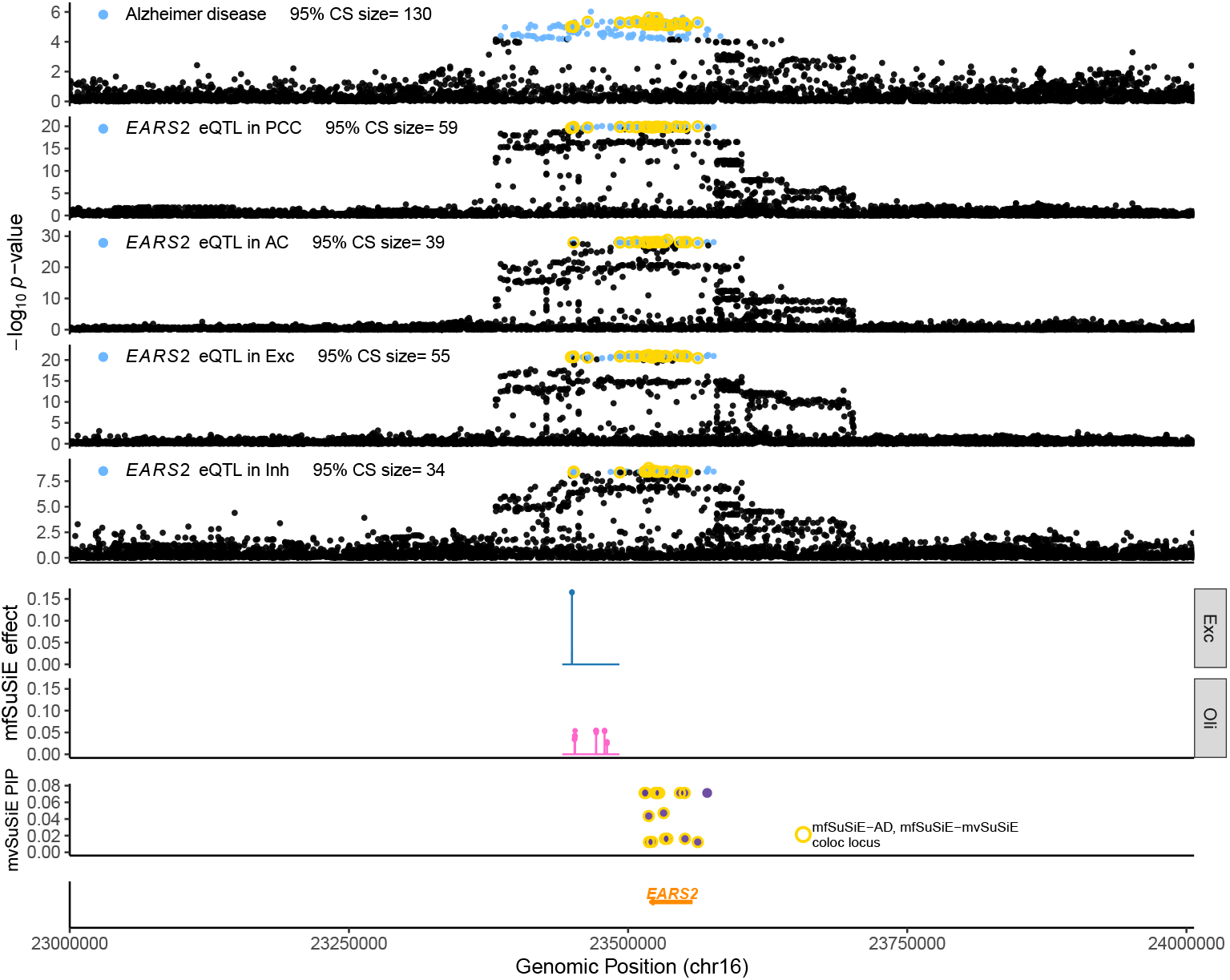
Multi-omic integration at *EARS2* AD risk locus. Top panels show single-trait SuSiE fine-mapping for AD GWAS and *EARS2* eQTLs across PCC, AC, Exc, and Inh. Blue points represent variants belonging to each trait’s 95% credible set. The lower panels summarize *mfSuSiE* fine-mapping of chromatin accessibility in excitatory neurons and oligodendrocytes. *mfSuSiE* identifies significant chromatin effects (LFSR < 0.01) in four cell types—Ast, Exc, Mic, and Oli. Similarly, *mvSuSiE* fine-mapping of *EARS2* expression detects significant eQTL effects (LFSR < 0.01) in bulk cortical tissues (AC, PCC, DLPFC) as well as in Exc, Inh, and Oli. Yellow circles mark variants supported across pairwise colocalization analyses connecting *mfSuSiE* (chromatin), *mvSuSiE* (expression), and AD GWAS, indicating that the shared variants fall within the same high-LD locus.

**Fig. 6.**
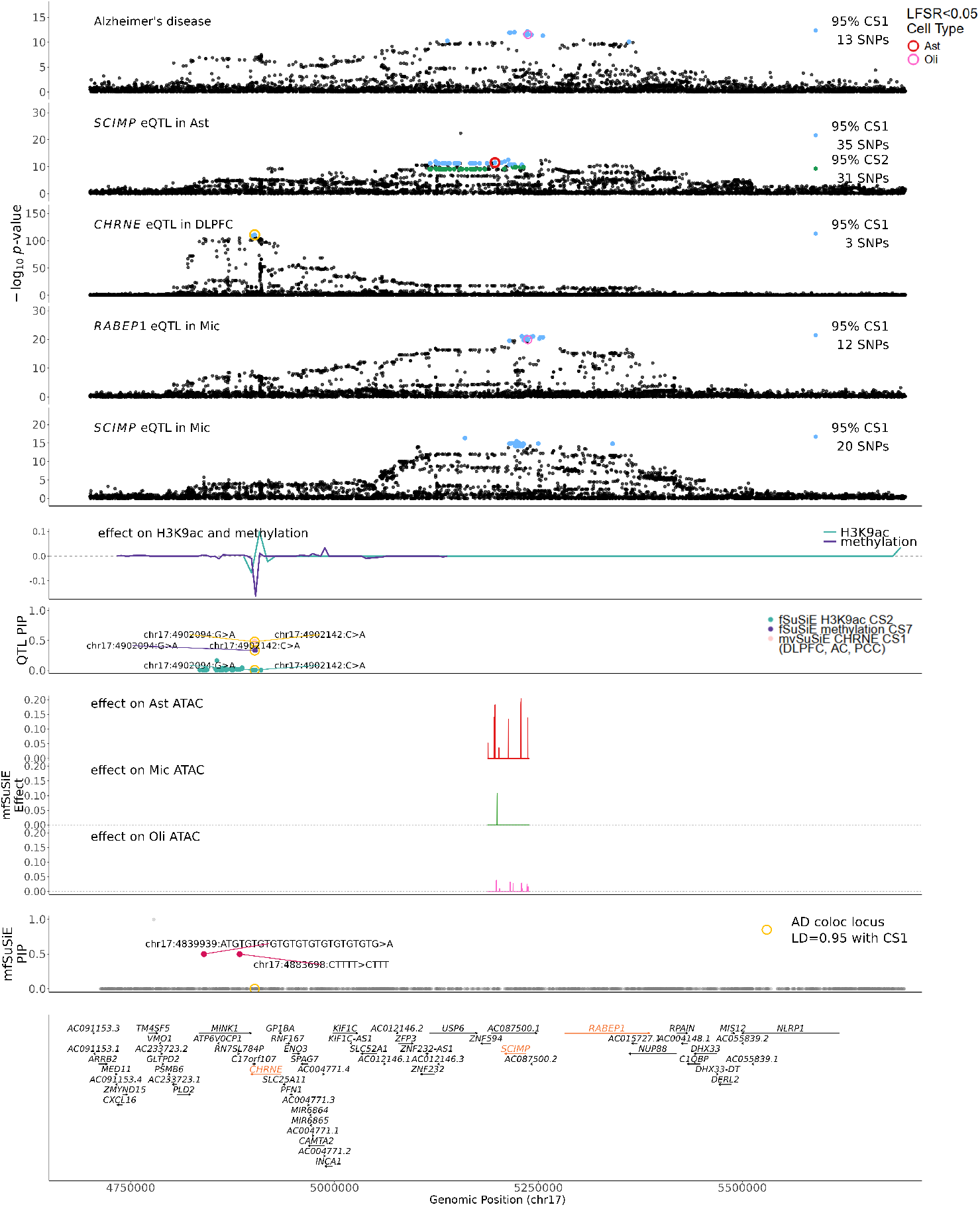
Multi-omic integration at the *CHRNE* /*SCIMP*/*RABEP1* AD risk locus. Integration of *mfSuSiE* chromatin accessibility fine-mapping with expression QTLs, epigenomic QTLs, and AD GWAS at chromosome 17: **(A)** Regional overview showing *mfSuSiE* –*SuSiE* colocalization analysis with AD GWAS at 4.9 Mb (PP.H4 = 0.86) and chromatin accessibility effects extending to the 5.2 Mb region across all six brain cell types. **(B)** *mvSuSiE* expression QTL fine-mapping at 5.2 Mb showing eQTL signals for *SCIMP* (microglia, astrocytes, excitatory neurons, inhibitory neurons, bulk tissues) and *RABEP1* (microglia, oligodendrocytes, bulk tissues), with target regions overlapping *mfSuSiE* chromatin accessibility effects. **(C)** *fSuSiE* fine-mapping of H3K9ac and DNA methylation QTLs at 4.9 Mb, with credible sets in high LD (*r*^2^ = 0.95) with *mfSuSiE* -identified variants. Effect directions are consistent with increased transcriptional activity: AD risk allele associates with increased H3K9ac and decreased DNA methylation. **(D)** Bulk tissue eQTL showing elevated *CHRNE* expression associated with AD risk variants at 4.9 Mb, supporting a regulatory cascade where variants modulate local epigenetic marks to control *CHRNE* expression while extending chromatin accessibility effects distally to regulate *SCIMP* and *RABEP1*.

To further investigate the 4.9 Mb region, we performed *mfSuSiE*-*mvSuSiE* colocalization to examine additional molecular QTL modalities. We observed bulk tissue eQTL effects at this position for *CHRNE* (anterior cingulate cortex, dorsolateral prefrontal cortex, and posterior cingulate cortex), but notably, no cell-type-specific eQTL signal. Moreover, while the caQTL effects extended distally to 5.2 Mb, the 4.9 Mb region itself showed no chromatin accessibility changes. To understand whether other epigenetic mechanisms might be operating at this locus, we applied *fSuSiE* (which can be viewed as a special case of *mfSuSiE* with a single phenotype) to histone modification and DNA methylation QTL data from FunGen-xQTL (details in **Methods**). Fine-mapping of H3K9ac (histone acetylation) and DNA methylation QTLs both identified credible sets at 4.9 Mb in high linkage disequilibrium (*r*^2^ = 0.95) with the *mfSuSiE*-identified caQTL. Effect directions were consistent with increased transcriptional activity: the AD risk allele associated with increased H3K9ac (transcription-promoting) and decreased DNA methylation (transcription-promoting) at regulatory elements near *CHRNE*, with corresponding elevated *CHRNE* expression in bulk tissue (Fig. 6).

Our analysis reveals regulatory architecture at this locus that potentially involves multiple coordinated mechanisms: AD risk variants at 4.9 Mb modulate local histone acetylation and DNA methylation to enhance *CHRNE* expression while simultaneously increasing chromatin accessibility across brain cell types, with effects extending distally to 5.2 Mb to control *SCIMP* and *RABEP1* expression. *CHRNE* encodes a subunit of nicotinic acetylcholine receptors critical for cholinergic signaling, with elevated expression linked to AD risk through inflammatory signaling and microglial activation. Both *SCIMP* and *RABEP1* have also been implicated in AD pathogenesis, with *SCIMP* involved in microglial immune responses and *RABEP1* in endosomal trafficking pathways relevant to amyloid processing. Notably, the 4.9 Mb and 5.2 Mb regions show only moderate linkage disequilibrium (*r*^2^ ≈ 0.13), suggesting they may act independently within a coordinated cis-regulatory network, with the top AD GWAS signal potentially reflecting aggregated LD effects across these regulatory elements. This multi-omic integration, enabled by *mfSuSiE*-based colocalization, allows us to clearly visualize and describe this complex regulatory architecture, which would be difficult to characterize without both the multi-cell-type fine-mapping framework and the integrated analysis across molecular modalities that *mfSuSiE* facilitates.

## DISCUSSION

We have introduced *mfSuSiE*, a method that extends functional fine-mapping to jointly analyze molecular QTLs across multiple cell types or contexts. Applied to single-nucleus chromatin accessibility data from aging brain, *mfSuSiE* substantially increased discovery of candidate regulatory variants compared to both conventional per-peak per-variant association testing and single-cell-type analysis with *fSuSiE*. Multi-cell-type analysis produced more credible sets, achieved higher fine-mapping resolution, and enabled characterization of extensive cell-type sharing patterns in chromatin accessibility regulation. An additional strength of *mfSuSiE* is its ability to accommodate phenotypes measured on partially overlapping or non-identical sample sets. Because the model can integrate molecular QTL data even when different modalities or cell types are profiled in distinct subsets of individuals, it substantially broadens the range of datasets that can be analyzed jointly. This flexibility makes *mfSuSiE* particularly well suited for large consortia studies, where multi-omic measurements are rarely obtained from the same donors. A key advantage of *mfSuSiE* beyond improved statistical power is that it enables systematic integration with other molecular modalities through colocalization analysis. By producing Bayesian inferences compatible with the broader *SuSiE* fine-mapping framework, *mfSuSiE* facilitates complex multi-omic integration pipelines that would otherwise require separate, disconnected analyses. As we demonstrated through 13-dimensional integration of chromatin accessibility, gene expression, and GWAS data, this capability allows researchers to trace regulatory effects across molecular layers and refine mechanistic understanding of disease-associated variants.

The framework we present is broadly applicable beyond chromatin accessibility QTLs. While we focused on single-nucleus ATAC-seq to illustrate multi-cell-type fine-mapping, *mfSuSiE* can be applied to any molecular QTL data that suits the functional regression framework, i.e., measurements are spatially structured along the genome and that the same genomic positions are measured across contexts. This includes analyzing histone modifications or DNA methylation across tissues or developmental stages, or combinations of multiple epigenetic marks measured in parallel. It can also be extended to analyzing any high-throughput sequencing data where coverage patterns represent the molecular phenotype. For instance, *mfSuSiE* could be applied directly to base-level RNA-seq coverage data instead of having to quantify gene-level expression first, which would enable joint fine-mapping of chromatin accessibility and gene expression from emerging multiome technologies. Such direct joint modeling would leverage the paired measurements more effectively than the post hoc colocalization approach we employed here.

Our work has several limitations representing directions for future research. The ROSMAP single-nucleus ATAC-seq data analyzed here, while demonstrating the advantages of multi-cell-type analysis, comprises a relatively small sample for systematic characterization of regulatory architecture in Alzheimer’s disease. We took a candidate region approach, guided by marginal association signals and established AD risk loci, rather than conducting genome-wide fine-mapping across all accessible regions. Applying *mfSuSiE* to larger single-nucleus chromatin accessibility datasets would enable more comprehensive discovery and better powered studies of cell-type-specific regulation in complex diseases. Second, the current implementation requires analyzing one molecular modality (e.g., chromatin accessibility) at a time within specified *cis*-regulatory windows around multiple molecular contexts (e.g., cell types). Future work could develop efficient implementation for simultaneously fine-mapping effects across multiple molecular layers within extended genomic windows, providing a more statistically principled and powerful framework for elucidating complex cis-regulatory networks.

Our work has several limitations that point to important directions for future research. Although the ROSMAP single-nucleus ATAC-seq dataset illustrates the advantages of multi-cell-type fine-mapping, it remains relatively small for a systematic analysis of regulatory architecture in Alzheimer’s disease. Because of this, we adopted a candidate-region strategy guided by marginal association signals and known AD risk loci, rather than performing genome-wide fine-mapping across all accessible regions. Applying *mfSuSiE* to larger single-nucleus chromatin accessibility cohorts will be essential for more comprehensive discovery and for better-powered characterization of cell-type-specific regulatory mechanisms in complex diseases. Additionally, although we analyzed six chromatin accessibility phenotypes jointly, our application focused on a single molecular modality. In principle, *mfSuSiE* could support simultaneous joint fine-mapping across heterogeneous molecular layers — ATAC-seq, DNA methylation, histone marks, RNA-seq coverage, and others — within extended genomic windows. We were unable to perform such analyses here due to data access constraints, relying instead on colocalization of Bayes factors across modalities. Future work will focus on performing joint fine-mapping of heterogeneous modality at a genome-wide scale.

## METHODS

### Simulation details

#### Simulated genotype

The fine-mapping regions for the simulations were selected uniformly at random from 94 breast cancer loci on autosomal chromosomes reported in [40] (see Supplementary Table 1 of that paper). The median size of a fine-mapping reagion was about 1 Mb. Similar to [41], we used sim1000G [28] to simulate genotypes of unrelated individuals based on the genotypes from the 1000 Genomes Phase 3 whole-genome sequencing [34]. First, we randomly selected a continent-of-origin label (EUR, AMR, AFR, EAS, SAS), then we simulated SNP genotypes using *N* = 100 individuals chosen uniformly at random from the 1000 Genomes samples with the selected continent-of-origin label. Within the fine-mapping region, we kept all biallelic SNPs with minor allele frequencies (MAFs) of 5% or greater; that is, SNPs in which the minor allele was observed at least 10 times out of the 2*N* = 200 chromosomes). For a single simulation, the genotype matrix, **X**, was a matrix with *N* = 100 rows (individuals) and *J* columns (SNPs), in which *J* ranged from approximation 1,500 to 4,000.

#### Simulated effects on molecular phenotypes

In these simulations, each modality of the molecular traits data were simulated from an *fSuSiE* model (more precisely, a multiple wavelet regression model with an SPS prior for the causal SNP effects). This involved the following steps: the effects of all causal SNPs *j* (on the wavelet scale) were simulated from an SPS prior, 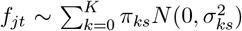, in which the first mixture component was always a “spike” at zero (*σ*_0*s*_ = 0); the effects of all non-causal SNPs *j* were set to zero, *f*_*jt*_ = 0; the effects in the measurement space were then obtained as **B** = **FW**^−1^; and then the molecular trait data, **Y**, were simulated from the multiple regression model (1), with 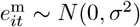. The residual variance *σ*^2^ was chosen so that **X** explained a specified total variance in **Y**.

Similar to [26] [42, 43], we simulated molecular trait data with different levels of smoothness: first, we drew a “smoothness parameter”, *µ*, uniformly at random between 0 and 1; then we set the *k* = 0 (“spike”) mixture component at each scale *s* to be *π*_0*s*_ = 1 − 2^−*µs*^. The intuition is that settings of *µ* closer to 1 produce “smoother” signals (larger effects at the largest scales), while settings of *µ* closer to 0 produce effects of similar size across all scales. For each simulations we select at random the number of causal variant from 1 to 20.

#### Univariate phenotypes

were simulated under standard linear model where the effect ***y***_*um*_ ∼ *Xβ* + ***e***_*um*_ where *um* stands for univariate trait number *m* and the entries of ***e***_*um*_ are independent *N* (0, *σ*^2^), causal SNP effect were sampled using a *β*_*j*_|SNP j causal ∼ *N* (0, 1).

### mfSuSiE model details

The *mfSuSiE* fitting procedure builds directly on the *fSuSiE* framework. At a high level, *mfSuSiE* fits a functional regression model—using the *fSuSiE* method [24]—to each data modality separately, and then combines the resulting evidence across modalities to form a joint Bayes factor for each SNP.

Below, we briefly summarize the single-modality functional regression procedure and then describe how evidence is integrated across modalities to fit a sum of single-effect priors.

### Multiple-trait wavelet regression

We begin by describing the core computation in *mfSuSiE*: the multiple-trait wavelet regression. To build intuition, we first consider the simpler case of a single trait.

#### Single-modality functional regression

Suppose we have *N* samples of a molecular trait measured at *T* spatial (or genomic) locations. Let **Y**_*m*_ ∈ ℝ^*N*×*T*^ denote the corresponding matrix of observations in modality *m*, with entries *y*_*it*_ for sample *i* ∈ {1, …, *N*} and location *t* ∈ {1, …, *T*} . Let ***x*** = (*x*_1_, …, *x*_*N*_)^⊺^ be a covariate of interest (e.g., a genotype).

We begin with the standard linear regression model

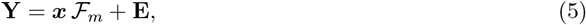

where ℱ_*m*_ = (ℱ_*m*1_, . . ., ℱ_*mT*_)^⊺^ is vector regression coefficients (“effects”), and **E** is the matrix of residuals in the measurement space.

#### Wavelet transformation

To exploit the decorrelating properties of wavelets, we transform the measurements to wavelet space via

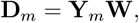

where **W** is the *T* × *T* orthogonal wavelet projection matrix. The transformed data matrix **D**_*m*_ ∈ ℝ^*N* ×*T*^ contains the wavelet coefficients (WCs). The orthogonality of **W** ensures invertibility and facilitates recovery of measurement-scale quantities from wavelet-scale estimates; see [24] for further background.

Applying the transformation to model (5) yields the wavelet regression model

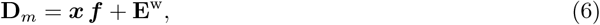

where ***f***_*m*_ = ℱ_*m*_**W** is the effect vector in wavelet space, and **E**^w^ contains the residuals in the wavelet space. We assume the residuals are independent across locations 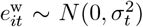 an assumption supported by the approximate “whitening” property of the wavelet transform [26, 27]. By contrast, the residuals in measurement space exhibit strong spatial correlation and cannot be assumed independent.

### Extending to multiple modalities

*mfSuSiE* assumes that (i) residual noise is independent across modalities, and (ii)effects across modalities are independent conditional on the causal SNP. Under these assumptions, fitting a multiple-modality wavelet regression amounts to: a) applying the single-modality wavelet regression independently to each of the *m* modalities, yielding a Bayes factor for the covariate ***x*** in each modality; and b) combining evidence by taking the product of the *m* modality-specific Bayes factors, which yields the *joint* Bayes factor for ***x***. This joint Bayes factor is then used within the *mfSuSiE* framework to fit the sum of single-effect priors across modalities.

### Single function for multiple-trait regression model

The “single function for multiple-trait regression” (SFMR) model is the functional counterpart of the “single function regression” (SFR) model [24]. The SFMR model is a multivariate multiple-trait wavelet regression model (6) with the constraint that exactly one of the covariates *j* ∈ {1, …, *J*} has a nonzero effect on the wavelet coefficients:

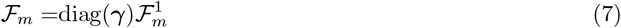

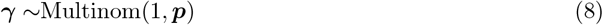

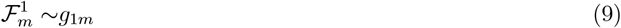

where 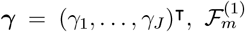 is a *J* × *T* matrix with entries 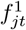, Multinom(*n*, ***p***) denotes the multinomial distribution with sample size *n* and multinomial probabilities ***p*** = (*p*_1_, …, *p*_*J*_), *g*_1*m*_ denotes the prior distribution for the effect in modality *m* (greater details are provided in the supplementary material), and diag(***γ***) denotes the *P* × *P* diagonal matrix in which the diagonal elements are given by the vector ***γ*** = (*γ*_1_, …, *γ*_*P*_).

Defined in this way, the SFMR model has the following properties: (i) ***γ*** is a binary vector in which exactly one of the elements is one and the rest are zeros; (ii) at most one of the rows of the ℱ_*m*_ contains nonzero values; and (iii) for each ℱ_*m*_ if there is a nonzero row in at least two matrix of coefficient (say ℱ_*m*1_ and ℱ_*m*2_) they have the same nonzero row, this corresponds to the fact that the SFMR models the effect of a single the causal variant at across all the modality.

### mfSuSiE model

The *mfSuSiE* model extends the SFMR model to allow at most *R* ≥ 1 nonzero effects. It is the multivariate functional counterpart to the functional Sum of Single Effects (*fSuSiE*) model, thus called the “mutlivariate functional Sum of Single Effects model” (*mfSuSiE*) model. It is a multiple-trait wavelet regression together with the following prior on **F**:

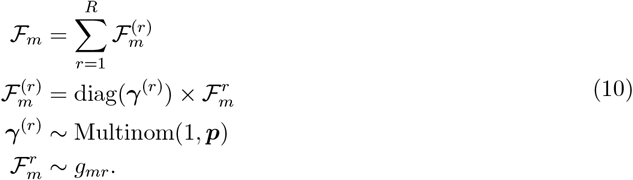

Since each binary vector 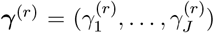 contains exactly one 1, by construction at most *R* rows of ℱ_*m*_ contain nonzeros. The *mfSuSiE* model reduces to the SFMR model when *R* = 1. Note that this model allows for a different prior on the effects *g*_*mr*_ for each effect *r* and modality *m* ; these priors are automatically adapted to the data using the algorithms described below.

### Posterior statistics

Here we define the key posterior quantities used in an *mfSuSiE* analysis (Fig. 4). We do not explain here how these quantities are computed; these details are given in the Supplementary Note.

As we briefly explained in the Methods overview, we have three main inference aims:

- *Variable selection:* identify the causal SNPs.
- *Modality annotation:* identify which modalities are affected by one or more SNPs.
- *Feature annotation:* identify the molecular features and locations that are affected by one or more SNPs in a given modality.

For variable selection, we compute *credible sets* (CSs): a CS is defined as a subset of {1, …, *J*} containing an effect SNP with high probability [3, 35]. More precisely, a level-*ρ* CS is defined as a set of SNPs that is as small as possible such that it has probability at least *ρ* of containing an effect SNP (a row of **F** containing at least one non-zero). The number of CSs should reflect the number of causal SNPs, and the size of a CS should reflect the number of plausible candidate effects SNPs. We calculate CSs as described in [3].

To determine which SNPs within a CS are the strongest candidates for being an effect SNP, we compute a *posterior inclusion probability* (PIP) for each SNP. The PIP for SNP *j* is defined as the posterior probability that at least one of the entries in the *j*th row of **f**_1_, …, **f**_*M*_ (noted **f**_1*j*_) is nonzero:

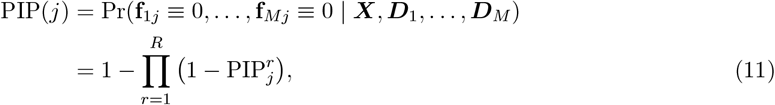

in which 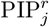 denotes the posterior probability that the *r*th single effect is nonzero for SNP *j*,

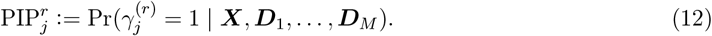

See Fig 4 for an illustration of PIPs and CSs.

The estimates of the SNPs on the molecular features in wavelet space are given by the posterior mean of 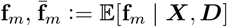. The SNP effects in measurement space are given by the posterior mean of 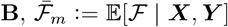. By elementary properties of expectations, this is simply 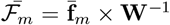, where **W**^−1^ denotes the inverse wavelet transform.

To identify affected features and locations, we compute 1 − *α* pointwise Bayesian credible intervals [36, 37] for elements *f*_*jt*_ and *b*_*jt*_ at selected SNPs *j*. We define “affected” as those elements in which the interval does not include zero. We refer to these intervals as “credible bands.” See Fig. 6 giving an example of the posterior means and the credible bands of *b*_*jt*_ for all epigenomic marks *t* = 1, …, *T* and for all sentinel SNPs *j*. The affected locations *t* are defined as the locations in which at least one of the credible bands for the sentinel SNPs at *t* does not contain a zero.

### Outline of an mfSuSiE analysis

Briefly, the minimal requirements for performing an *mfSuSiE* analysis are:

i) An series *m* of *N* × *T*_*m*_ matrices, each containing *N* molecular trait measurements at *T*_*m*_ locations, some rows can contain missing data.
ii) An *N* × *J* matrix, 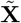, containing the genotype information for the *J* SNPs to be fine-mapped after removing the linear effects of the selected covariates.
iii) *R*^max^, an upper limit on *R*, the number of single functions in the *fSuSiE* model. Unless specifically mentioned, we set *R*^max^ = 10.

An additional input is optional but recommended:

v) The positions (e.g., base-pair positions) for each modality 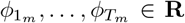 corresponding to the locations 1, …, *T* . If not specified, the positions are assumed to be evenly spaced. Molecular traits are typically not measured at evenly spaced positions, so providing this information will produce more accurate results.

Beyond this, the *mfSuSiE* software includes many other settings and tuning parameters which may be adjusted as needed.

The basic steps of an *mfSuSiE* analysis are as follows:

1. *Compute the wavelet coefficients*. Compute the WCs for each modality, **D**_*m*_, from the molecular trait data, **Y**_*m*_. For computational efficiency, **D**_*m*_ is obtained using the standard discrete wavelet transform (DWT). (In later steps, a second set of WCs are computed using the translation-invariant wavelet transform to improve accuracy.) For all the results presented in this paper, we used the (undecimated) wavelet transform with Daubechies least-asymmetric orthonormal compactly supported wavelets and with 10 vanishing moments (see Chapter 2 of [44]). Note that the *fSuSiE* software currently supports any wavelet transform that is implemented in the wavethresh R package [44].
2. *Search for a good upper limit on R (optional). R* determines the number of single functions and, correspondingly, the number of causal SNPs. If *R* is too small, *mfSuSiE* may miss some causal SNPs; on the other hand, if *R* is too large, *mfSuSiE* may take a long time to run. To reduce the overall compuational speedd we implemented the same *ad hoc* procedure to find a reasonable initial estimate of *R* as in *fSuSiE*. By default, this procedure starts by fitting an *mfSuSiE* model with *R* = 3. If all *R* = 3 single functions are kept, *R* is increased by one. This procedure iterates until one or more single functions are pruned, or if the upper limit, *R*^max^, is reached.
3. *Fit mfSuSiE model*. We use an iterative algorithm, described below, to fit the *mfSuSiE* model to the data **X, D**. This includes estimating the priors 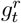 and the residual variance *σ*^2^. During estimation of the priors, some single functions may be pruned, and therefore *R* is an upper bound on the final number of single functions (and CSs).
4. *Compute SNP-level posterior quantities: CSs and PIPs*. Compute CSs and PIPs from the *fSuSiE* model fitted in the previous step.
5. *Filter credible sets (optional)*. One may filter out the CSs with low “purity” (purity is defined as the smallest absolute correlation among all pairs of SNPs in the CS). This often improves quality or interpretability of the fine-mapping results.
6. *Compute posterior effect estimates and credible bands*. The final step is to compute posterior mean estimates of *b*_*jt*_, and corresponding 1 − *α* pointwise credible bands, *α* ∈ [0, 1], for each location *t* and sentinel SNP *j* (the SNP with the largest PIP in each CS). To address possible inaccuracies with the DWT (see Chapter 9 of [27]), we compute a new WC matrix 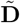 using the translation-invariant wavelet transform (TIWT), also known as the stationary wavelet transform. (In brief, the TIWT modifies the original DWT by applying it to shifted copies of the signal, then the WCs are averaged across the shifted copies.) Since the TIWT greatly increases computational effort, we use this TIWT only in this final step.

### Molecular QTL data resource and processing

We analyzed *n* = 92 ROSMAP snATAC-seq fragment files obtained from dorsolateral prefrontal cortex (DLPFC). Raw BCLs were demultiplexed using `cellranger-atac mkfastq`, aligned to GRCh38 with `cellranger-atac count`, and processed using ArchR with stringent QC. High-quality nuclei were defined as having TSS enrichment *>* 6 and 1,000–100,000 unique fragments, with doublets removed at sample and cluster levels. Cell types were assigned by marker-gene accessibility, producing six major lineages: Astrocytes (Ast), Excitatory neurons (Exc), Inhibitory neurons (Inh), Microglia (Mic), Oligodendrocytes (Oli), and OPCs [29].

Fragment files were aggregated to form cell-type–specific donor-level pseudobulk profiles using a fixed 500-bp peak set (363,775 peaks). Peaks overlapping hg38 blacklist regions were removed, leaving 363,746 peaks. Donors with *>* 50 nuclei per cell type were retained, yielding a total of 84 unique donors across all cell types. We filtered low-coverage peaks, normalized counts, and generated technical covariate-adjusted residuals. Technical covariates included sequencing batch, post-mortem interval (PMI), log(nuclei count), TSS enrichment, nucleosome signal, total fragments, percent reads in peaks, median peak width, and log(unique peaks).

#### Identification of regions with potential shared effect

We mapped chromatin accessibility QTLs (caQTLs) across six cell types using TensorQTL. For each pseudobulk peak, SNPs within a *±*1 Mb window centered on the peak midpoint were tested. Covariates included age at death, sex, genotype, and phenotype principal components (PCs). Following [29], we retained only SNP–peak associations with minor allele count (MAC) *>* 5 and permutation-based empirical p-value using the Beta approximation *p*_beta_ < 0.005. These SNP–peak pairs were defined as “gPeaks.” To ensure multi-cell-type relevance, we selected only gPeaks significant in ≥ 2 cell types, resulting in ≥ 12 multi-cell-type caQTL regions. Each gPeak (500 bp) was expanded symmetrically to a 51.2 kb window, which forms the first source of candidate regions for *mfSuSiE*. Results of these analysis is summarized in Table 1.

**Table 1.** Chromatin accessibility QTL association testing summary by cell type.

| Cell Type | Sample Size | Peaks | gPeaks<sup>†</sup> | Shared gPeaks<sup>‡</sup> |
| --- | --- | --- | --- | --- |
| Astrocytes (Ast) | 71 | 111,325 | 422 | 2 |
| Microglia (Mic) | 70 | 74,210 | 285 | 0 |
| Excitatory neurons (Exc) | 61 | 201,651 | 766 | 5 |
| Inhibitory neurons (Inh) | 75 | 127,504 | 528 | 7 |
| Oligodendrocytes (Oli) | 83 | 176,039 | 1,198 | 10 |
| Oligodendrocyte precursor cells (OPC) | 73 | 66,154 | 227 | 1 |

<sup>†</sup>gPeaks correspond to peaks with at least one SNP reaching $p_\beta < 0.005$ . <sup>‡</sup>Shared gPeaks are peaks with at least one SNP reaching $p_\beta < 0.005$ in at least two cell types.

To complement caQTL-driven regions, we incorporated *N* = 156 Alzheimer’s disease GWAS loci. For each locus, the top SNP was identified from AD summary statistics and expanded by *±*25.6 kb to create a 51.2 kb region. This produced 156 AD-associated candidate regions, which were combined with the 12 caQTL regions to yield a total of 168 fine-mapping regions. Only SNPs with MAF *>* 0.05 in ROSMAP WGS were retained for downstream modeling.

For every selected 51.2 kb region, we calculated base-level coverage directly from donor fragment files. Coverage was aggregated into 50-bp bins, generating *T* = 1024 genomic positions (51.2 kb / 50 bp), *M* = 6 cell types, and *N* = union of QC-passed donors across the 6 cell types (up to *n* = 84). This resulted in an *N* × *T* matrix per cell type, and an *N* × *T* × 6 multivariate phenotype tensor per region. These phenotypes match the structure illustrated in Fig. 1 (51.2 kb windows, binned base-level coverage).

For each 51.2 kb region, we extracted all SNPs within a *±*500 kb (total 1 Mb) window centered on the region midpoint. This window typically contains ∼ 2, 000–4,000 SNPs per region. Identical covariates to those used in TensorQTL were applied in *mfSuSiE*, including sequencing batch, PMI, age at death, sex, genotype PCs, and technical QC metrics (TSS enrichment, nucleosome signal, log(nuclei), etc.). The final *mfSuSiE* input therefore consists of phenotype data as 51.2-kb region × 1024 bins × 6 cell types *×N* donors, genotype matrix restricted to the 1 Mb window per region, covariates identical to TensorQTL, and variants with MAF *>* 0.05.

## Data availability

The datasets analyzed, including the ADSP R4 WGS genotype data, are available for application and download from the NIAGADS Data Sharing Service, https://dss.niagads.org. The code implementing the data processing and analysis pipelines for is available at https://statfungen.github.io/xqtl-protocol. The complete set of QTL data and QTL-GWAS integration models will be made publicly available at https://synapse.org prior to publication for registered Synapse users, as per the Data Management and Sharing policy of the FunGen-xQTL project. Other data sets used include: 1000 Genomes Phase 3 whole-genome sequencing data, https://ftp.1000genomes.ebi.ac.uk/vol1/ftp/release/20130502/; GENCODE Ensembl/Havana database, https://useast.ensembl.org/info/genome/genebuild/annotation_merge.html;

## Code availability

The mfsusieR R package is available on GitHub at https://github.com/stephenslab/mfsusieR (3-clause BSD license). PLINK 1.9 (https://www.cog-genomics.org/plink/); TensorQTL 1.0.8 (https://github.com/broadinstitute/tensorqtl/); sim1000G 1.40 (https://github.com/adimitromanolakis/sim1000G); SeqSIMLA (https://seqsimla.sourceforge.net/); susieR 0.14.7 (https://github.com/stephenslab/susieR/); coloc 5.2.3 (https://github.com/chr1swallace/coloc/); ashr 2.2-63 (https://github.com/stephens999/ashr/); mixsqp 0.3.18 (https://github.com/stephenslab/mixsqp/); wavethresh 4.7.2 (https://cran.r-project.org/package=wavethresh); R 4.3.3 (https://www.r-project.org).

## Acknowledgments

We thank the staff at the Research Computing Center at the University of Chicago for providing the high-performance computing resources used to implement the numerical experiments. We thank Angela Helfrich and Mark Bronnimann from Amazon Web Services for providing cloud computing support for real-world data analysis. We also thank the members of the Alzheimer’s Disease Sequencing Project Functional Genomics Consortium (FunGen-AD) for providing the FunGen-xQTL resource. This work was supported in part by NIH grants R01AG076901 and R01AG086467 (to G.W., A.L.), U01AF072572 (to P.L.D.) and a grant from the Urbut Family Foundation (to G.W.). This project is supported by the Eric and Wendy Schmidt AI in Science Postdoctoral Fellowship, a Schmidt Sciences, LLC program. Additional support came from the University of Chicago Data Science Institute through the 2024 AI+Science Research Initiative. This research was conducted using data from the Religious Orders Study and the Rush Memory and Aging Project (ROSMAP). We thank the participants and investigators of these studies.

## Author contributions

GW and WD jointly supervised research. WD developed the method and algorithm with input from AL and GW. GW and WD conceived and designed the experiments and data analysis. WD implemented the methods comparisons in simulations with input from AL. AL conducted data applications and interpreted the results with input from WD and GW. PLD contributed and supervised molecular QTL data production from the ROSMAP cohort. The Alzheimer’s Disease Functional Genomics Consortium contributed additional data resources. WD, AL and GW wrote and revised the manuscript. All authors critically reviewed the manuscript, suggested revisions as needed, and approved the final version.

## Supplementary figures

**Supplementary Figure 5.**
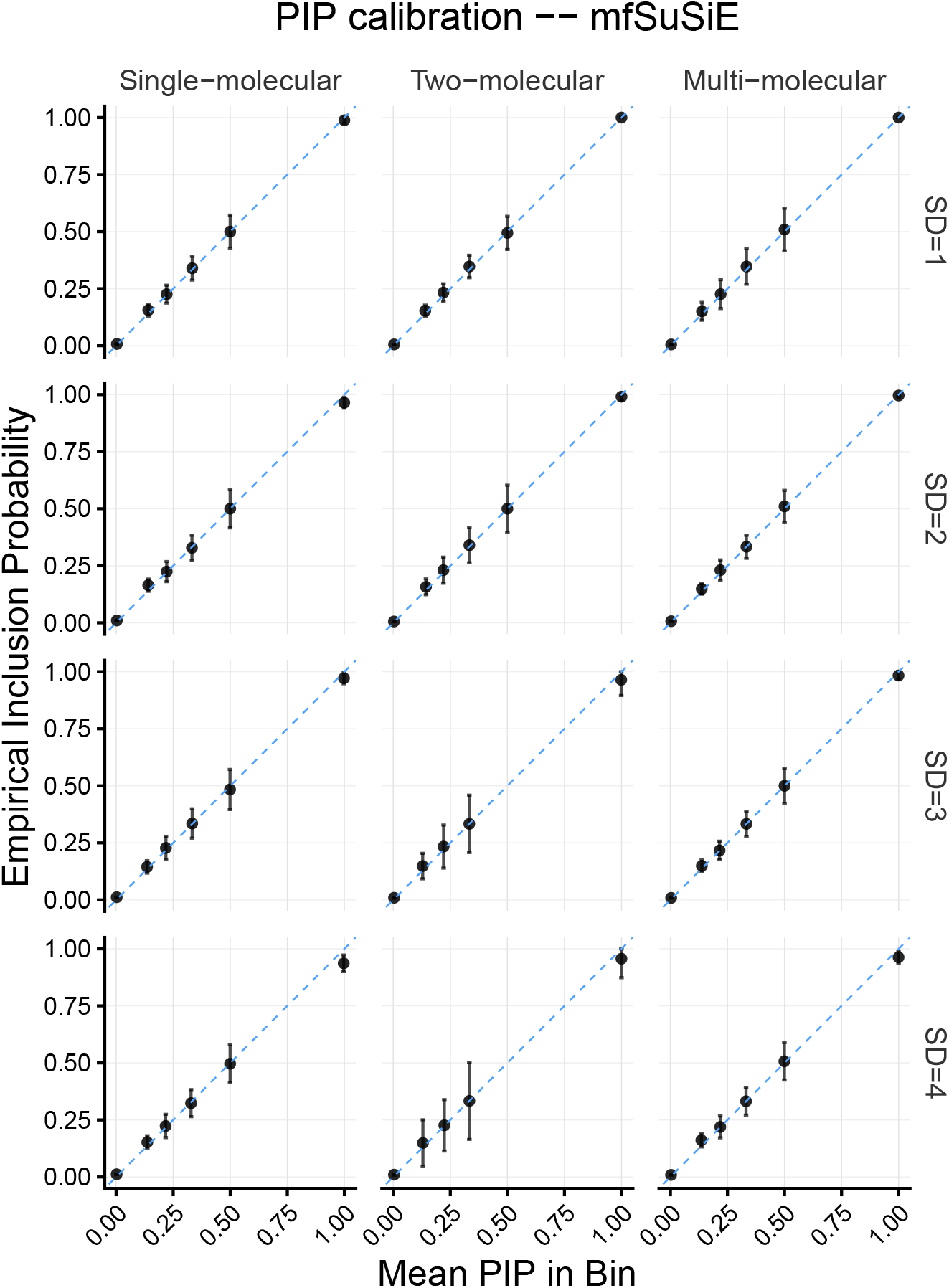
PIP calibration assessment for *mfSuSiE* method. Posterior inclusion probability calibration evaluation for the *mfSuSiE* method across three modeling scenarios under noise levels ranging from SD = 1 to 4: (1) Single molecular phenotype (32 columns); (2) *mfSuSiE* with 2 molecular phenotypes (32 + 64 columns); (3) Full model with *mfSuSiE* using 2 molecular phenotypes + 3 univariate phenotypes. The calibration plots assess whether reported posterior inclusion probabilities match empirical frequencies, demonstrating the reliability of uncertainty quantification across different noise conditions and model complexities.

**Supplementary Figure 6.**
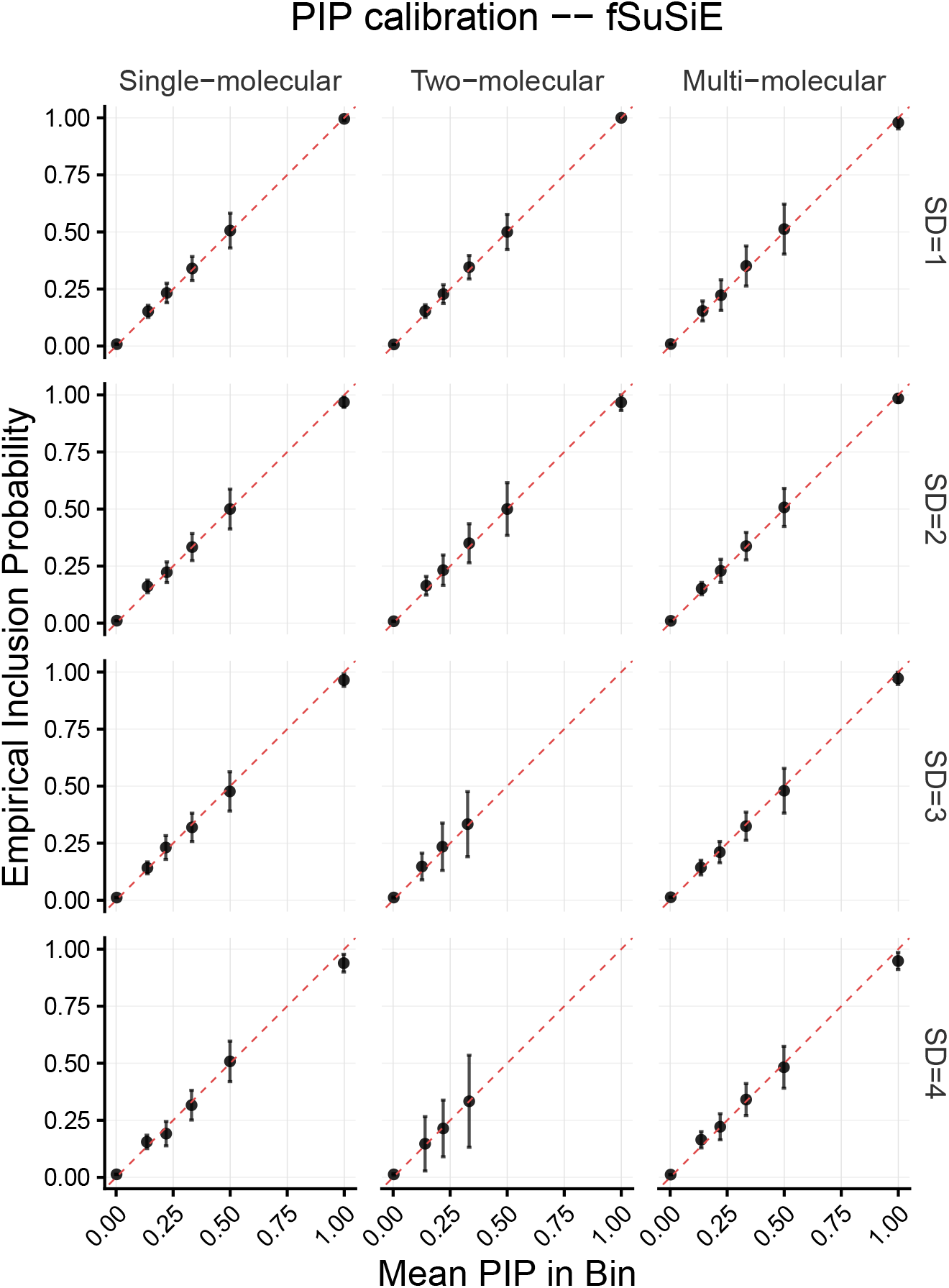
PIP calibration assessment for *fSuSiE* method. Posterior inclusion probability calibration evaluation for the single-cell-type *fSuSiE* method across three modeling scenarios under noise levels ranging from SD = 1 to 4: (1) Single molecular phenotype (32 columns); (2) *fSuSiE* with single molecular phenotype (32 columns) compared against *mfSuSiE* with 2 molecular phenotypes; (3) *fSuSiE* using only the first molecular phenotype compared against full model *mfSuSiE* . The calibration plots assess whether reported posterior inclusion probabilities match empirical frequencies, enabling direct comparison of calibration quality with the *mfSuSiE* method and demonstrating uncertainty quantification performance across different noise conditions.

**Supplementary Figure 3.**
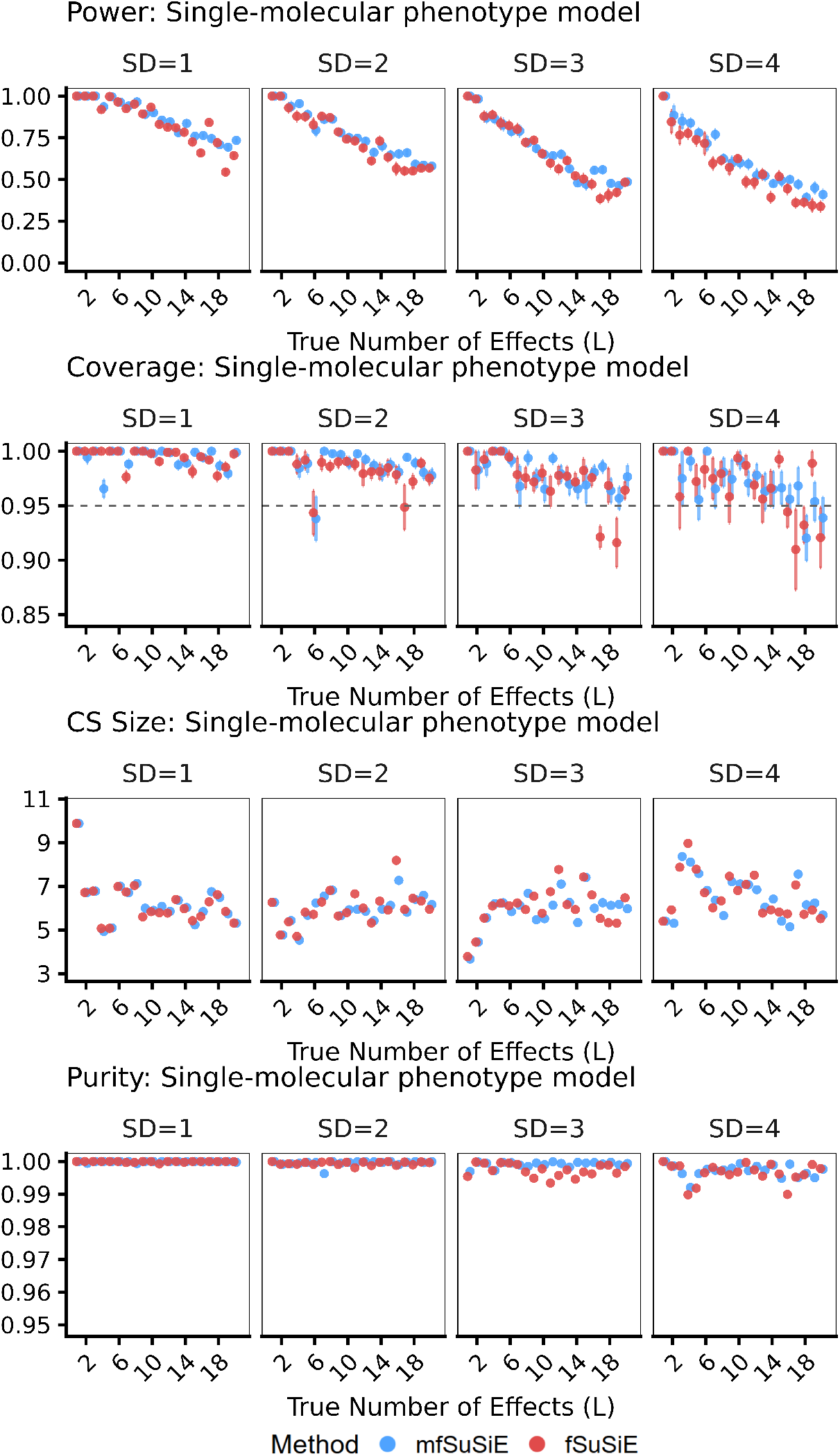
Complete simulation results for single molecular phenotype model. Comprehensive performance comparison between *mfSuSiE* and *fSuSiE* across four noise levels (SD = 1, 2, 3, 4) for the single molecular phenotype scenario. Both methods analyze the same single molecular phenotypes with 32 columns. Performance metrics include power (proportion of true causal SNPs included in at least one credible set), coverage (proportion of credible sets containing a true causal SNP), credible set size distribution, and purity (proportion of causal SNPs within each credible set). Each panel shows results across different simulation parameters with n = 100 samples.

**Supplementary Figure 7.**
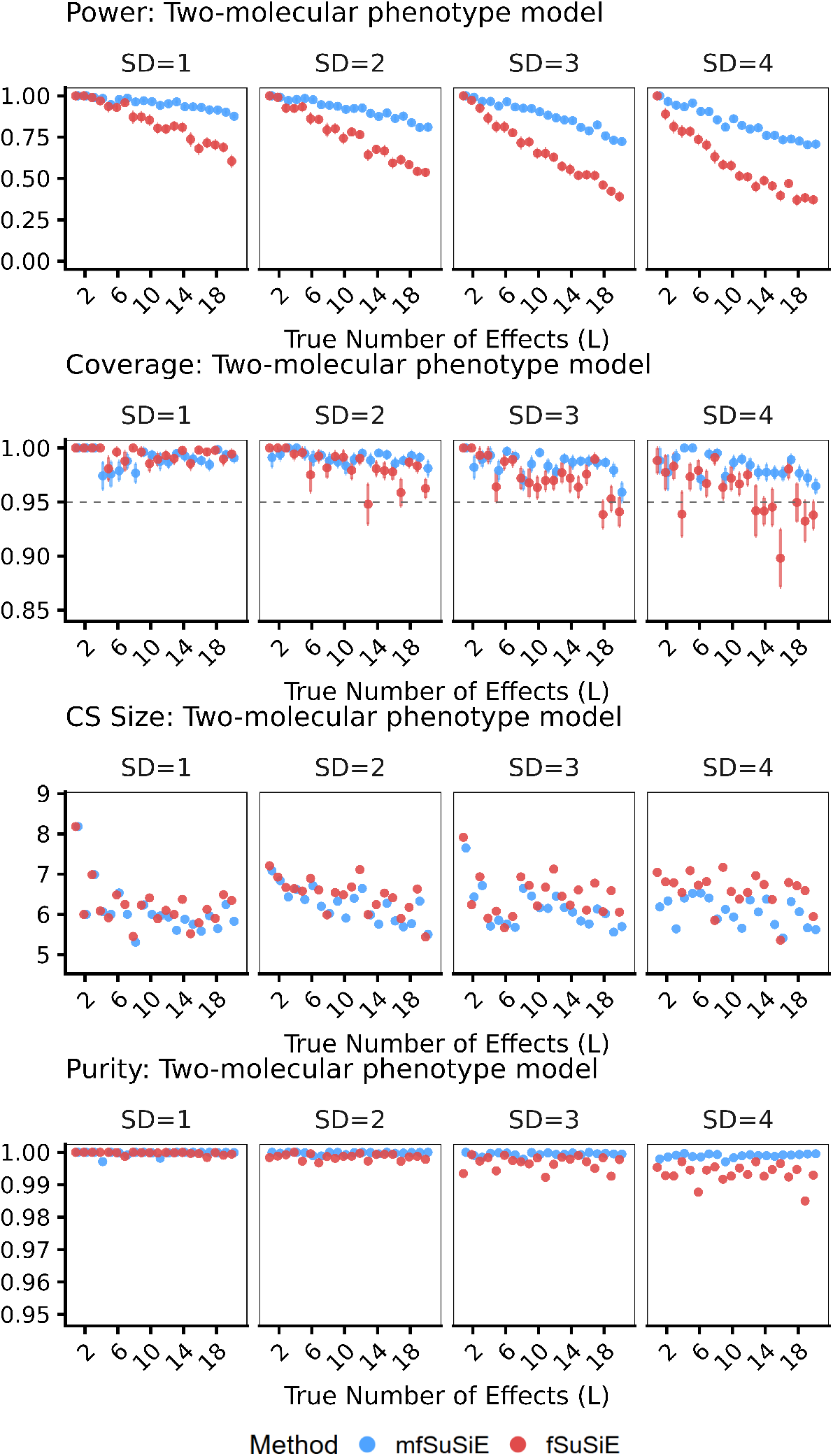
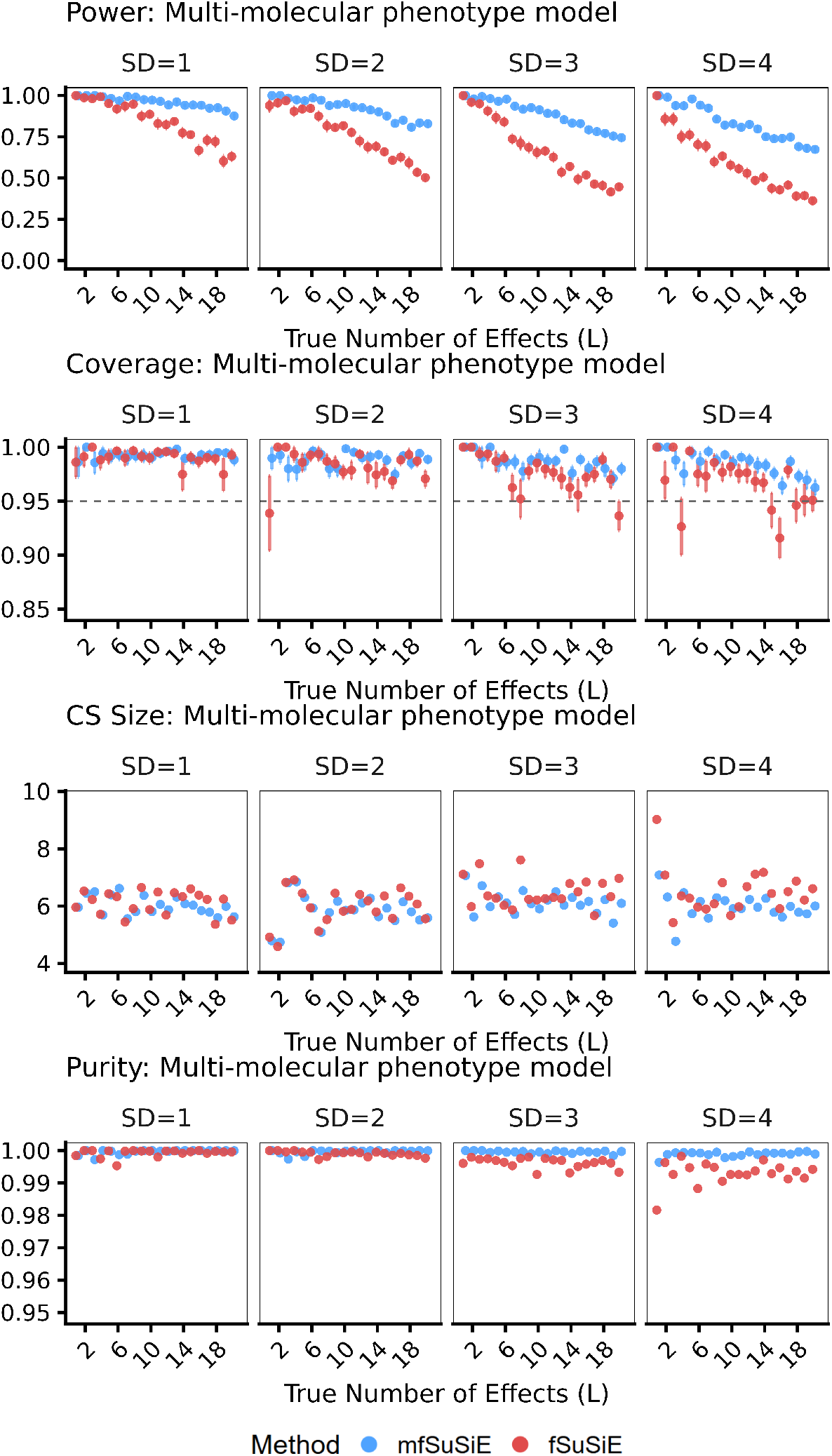
Complete simulation results for two molecular phenotypes model. Comprehensive performance comparison between *mfSuSiE* and *fSuSiE* across four noise levels (SD = 1, 2, 3, 4) for the two molecular phenotypes scenario. *mfSuSiE* analyzes both molecular phenotypes (32 + 64 columns) jointly, while *fSuSiE* analyzes only the first molecular phenotype (32 columns) independently. This comparison demonstrates the advantage of joint modeling when multiple related molecular phenotypes are available. Performance metrics include power, coverage, credible set size distribution, and purity. Each panel shows results across different simulation parameters with n = 100 samples.

**Supplementary Figure 8.**
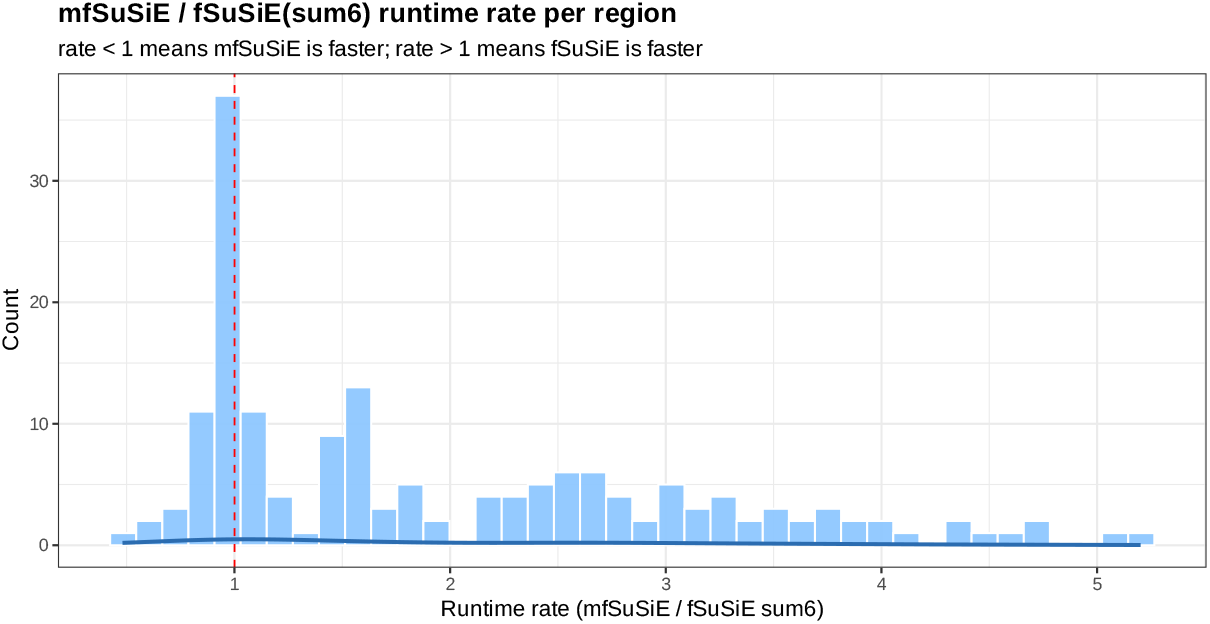
Runtime performance of *mfSuSiE* demonstrates fast execution and linear scaling with number of cells. *mfSuSiE* runs efficiently and scales roughly linearly with the number of cells analyzed. The analysis shows consistent performance across different dataset sizes, making it suitable for large-scale single-cell applications. Results are shown for credible sets with purity ≥ 0.8 and PIP ≥ 0.1.

**Supplementary Figure 9.**
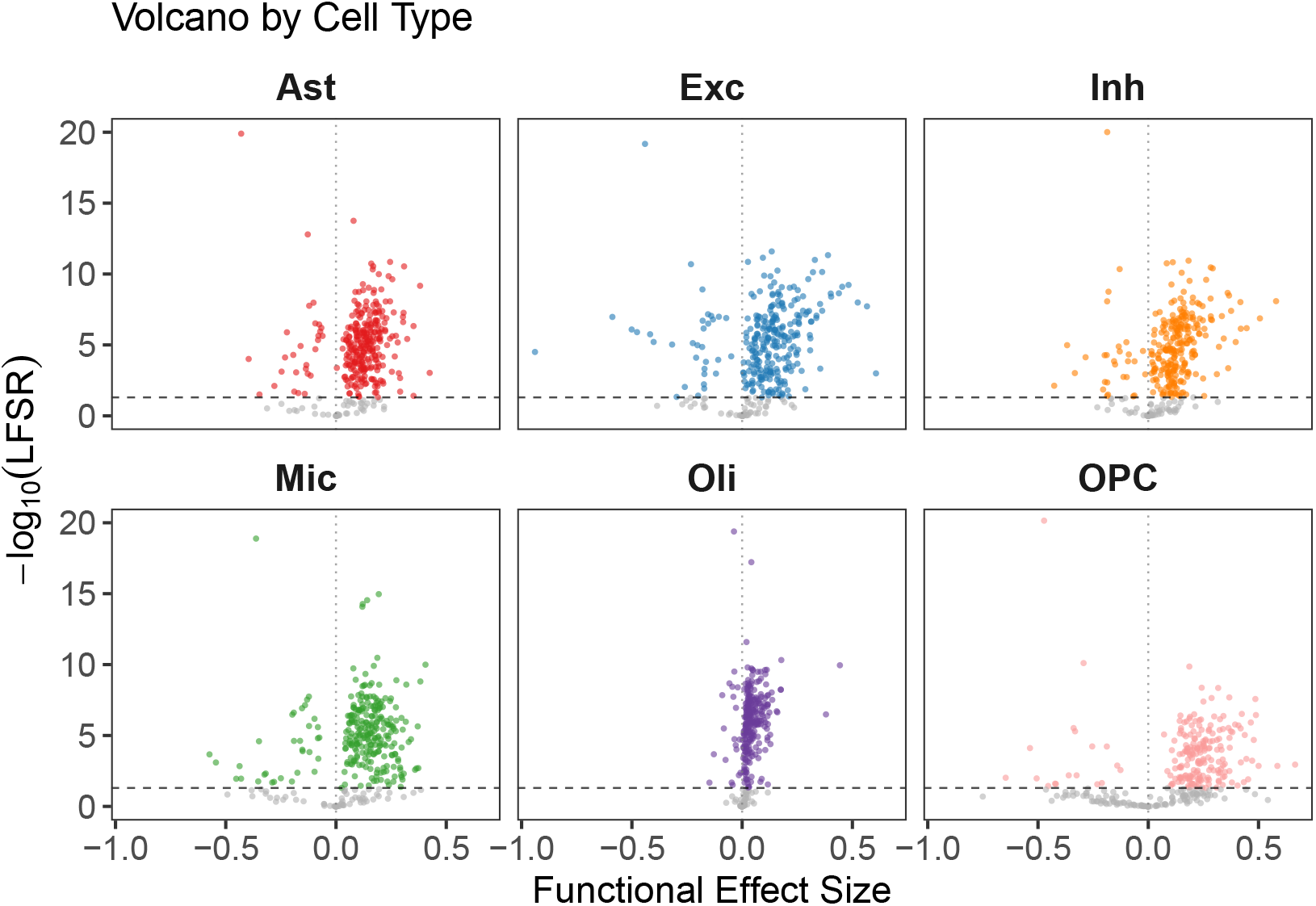
Cell-type-specific volcano plots of *mfSuSiE* fine-mapping results. Each panel displays the effect size (x-axis) versus −log_10_(LFSR) (y-axis) for variants within credible sets (CSs) for a given cell type. Local false sign rate (LFSR) represents the posterior probability of incorrect effect direction; lower values indicate higher confidence. Variants are included if they belong to CSs with purity *>* 0.8 or have posterior inclusion probability (PIP) *>* 0.1 outside 95% CSs. Dashed horizontal line indicates LFSR = 0.01 significance threshold. The y-axis is truncated at 20 for visualization.

**Supplementary Figure 10.**
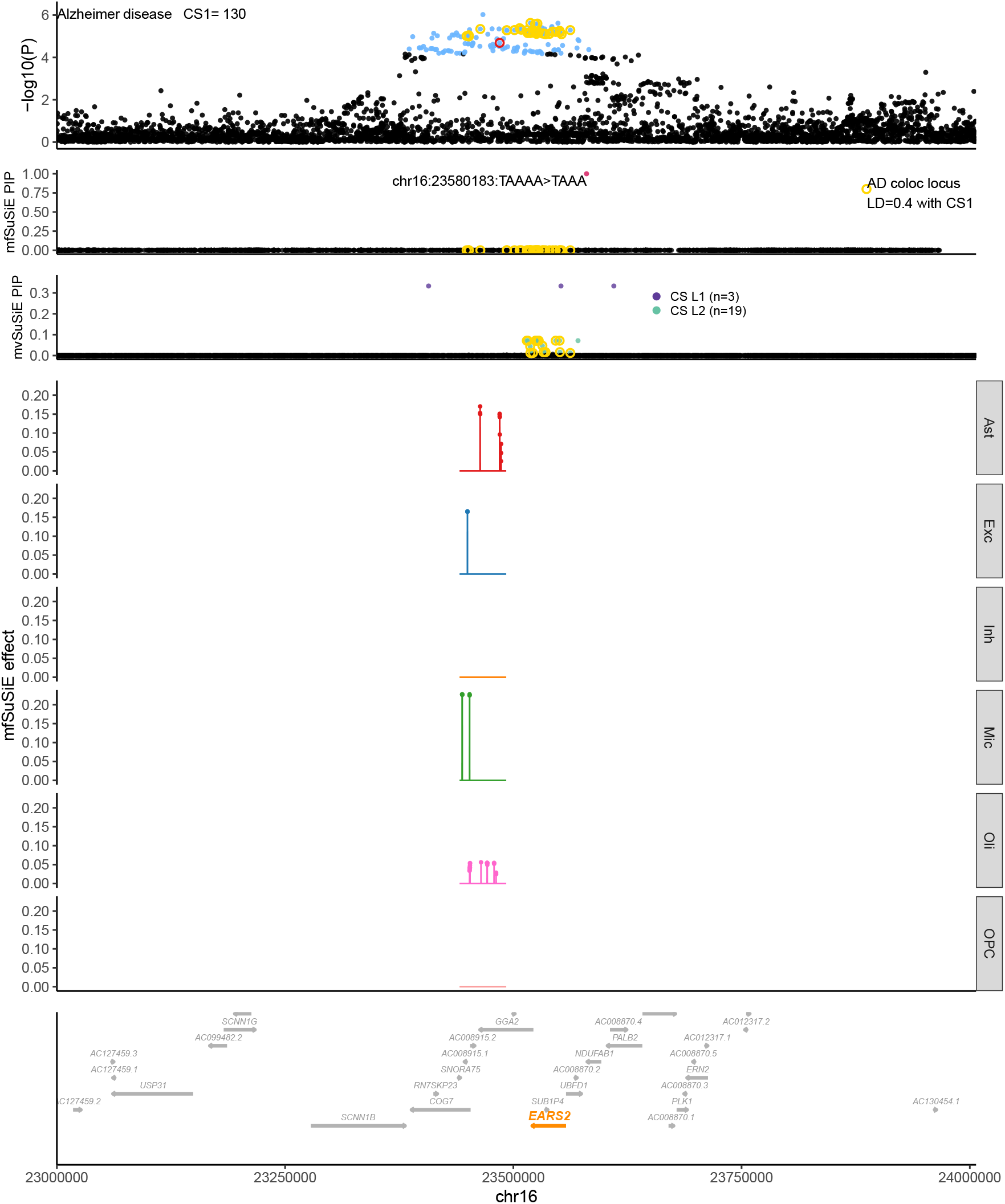
EARS2 case study.

## Supplementary text

### Notation

Matrices are denoted by bold, uppercase letters (e.g., ***A***). Three-dimensional tensors are denoted by bold, uppercase letters with three dots as indexes (e.g., ***A***_.,.,._). Furthermore, ***A***_.,*i,j*_ corresponds to the vector in the tensor ***A***_.,.,._ when fixing the two other dimensions. Similarly, ***A***_.,.,*j*_ corresponds to the matrix in the tensor ***A***_.,.,._ when fixing one dimension. Column vectors are denoted by bold, lowercase letters (***a***), and scalars are denoted by letters (*a* or *A*). For indexing, we use capital letters to denote the total number of elements and the corresponding lowercase symbol to denote the index, e.g., *i* = 1, …, *I*. For wavelets, we use two sets of indexes; when necessary, we use the classic double indexing of wavelet [27], where the index *s* ∈ 0, …, *S* corresponds to the scale (or level of resolution) index and *l* ∈ 1, …, 2^*s*^ + 1 to the location index. When possible, we simply index wavelet coefficients by *t* ∈ 1, …, *T* . We denote sets by calligraphic letters (e.g., 𝒜) and tensors are denoted using the “frak” font (i.e. 𝔄𝔅ℭ𝔇𝔈𝔉). To lighten the notation, collections of elements described by a set of indexes are indicated between parenthesis without specifying the index support; e.g. (*a*_*u,h*_) = (*a*_*u,h*_)_*u*∈𝒰,*h*∈ℋ_

## 1. Inference in the wavelet space details

When applying the wavelet transform by the right, and noting **D** = **YW** the empirical wavelet coefficient, **F**_*j*_ = ℱ_*j*_**W** the wavelet transform of the *j* function and **U** = **U**^*′*^**W**, where the entries of **U** are iid *N* (0, *σ*^2^). The previous model can be written as

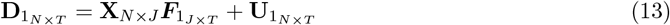

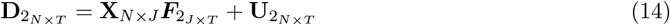

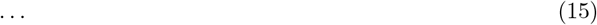

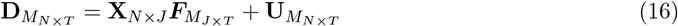

For the sake of simplicity, we assume that each **D**_*m*_ and **X** is scaled and center and so the intercept for each modality to be null. We also use the classic double indexing of wavelet coefficient (s,l) where *s* ∈ [1, log_2_(*T*)] and *l* ∈ [0, *s* − 1] instead of the *t* ∈ [1 : *T*] notation.

### Likelihood

We assume that the components of transformed data D to be independent.

Then the likelihood of D can be factorised as

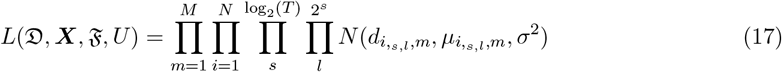

Where 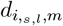 is the observed wavelet coefficient *s, l* for individual *i* in frame *m* and 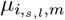is mean of wavelet coefficient *s, l* for individual *i* in frame *m*, .

### Prior

Let’s denote 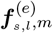 the p-vector corresponding to the *e*^*th*^ effect of each SNP for wavelet coefficient *s, l* in modality *m* (i.e., 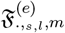). We model the effect of the SNPs on the different modalities through the following sum of single effect model

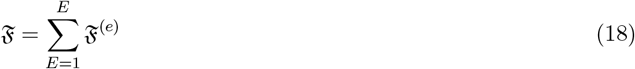

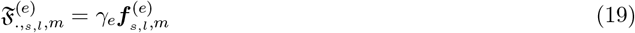

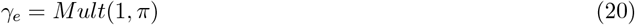

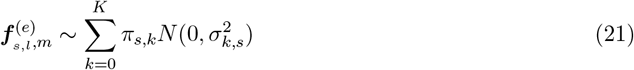

For the sake of lightening the notation we assume that *σ* in equation 17 and the grid of *σ*_1:*K*_ is the same for every scale and every function; in practice, we use a different grid and different noise level of for each scale and function. We refer to the model described in equation 18 as the (sum) of multiple functions model (*mfSuSiE* for short)

## 2. Fitting the sum of multiple functions Bayesian regression

First we start by detailing how to fit the following basic model

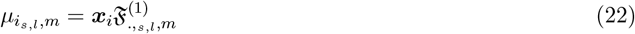

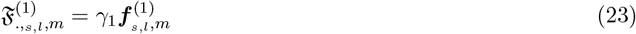

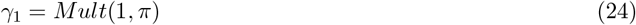

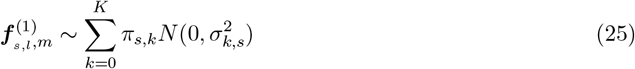

which we refer to as the single effect multivariate function Bayesian regression model. This fitting procedure will be used afterward to fit model *mfSuSiE* 18.

### 2.1. Remainder of the univariate Bayesian regression

Let’s consider the following simple Bayesian functional regression

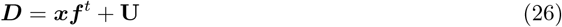

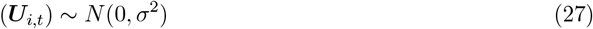

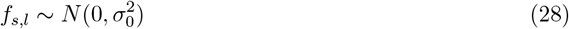

Where ***D*** is a matrix of wavelet coefficients of dimension *N* × *T*, ***x*** is a vector of length *N*, ***f*** is a vector of length *T*, ***U*** is a matrix of dimension *N* × *T*, and *i* ∈ [1 : *N*] and *t* ∈ [1 : *T*]. Then by assuming independence of the wavelet coefficients, the component of ***f*** can be estimated via maximum likelihood, as well as their corresponding variance 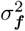are:

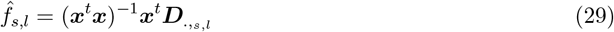

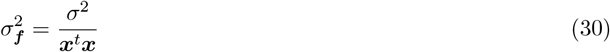

Where 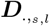 is the column of ***D*** corresponding to the empirical wavelet coefficients *s, l*. Thus, the posterior distribution of each component of ***f*** is

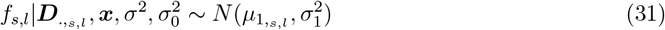

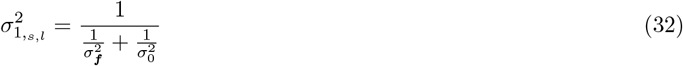

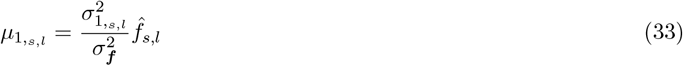

Following [38], Bayes factor for comparing the estimate ***f*** by the simple Bayesian functional regression with the null (*H*_0_ : ***f*** ≡ 0) is given by:

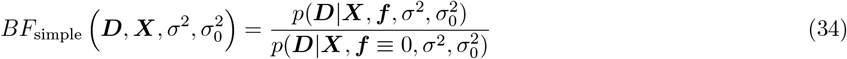

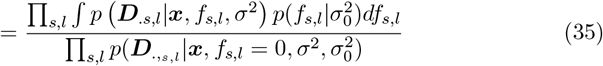

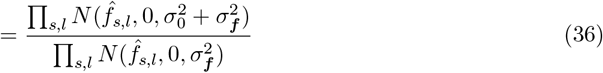

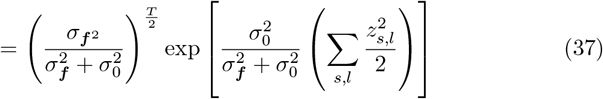

Where 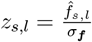 .

### 2.2. Fitting single effect multivariate function Bayesian regression model with mixture prior using a single covariate

Here we fit model 22 in the case where we assume that ***X*** as a single column (i.e., ***X*** = ***x*** and so *γ*_1_ = 1). Suppose that 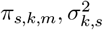 are given. Let’s introduce some latent state each *s, l, m*

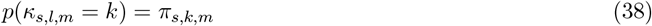

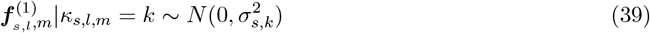

Using results from equations 32 and 33, the posterior distribution of *f*_*s,l*_ given that *κ*_*s,l*_ = *k* is

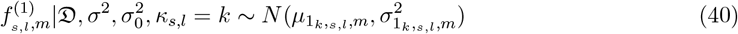

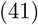

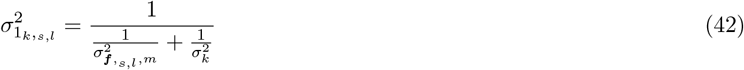

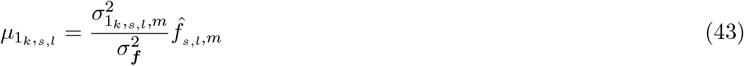

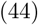

Thus the posterior mean of ***f*** is given by

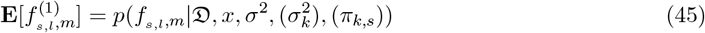

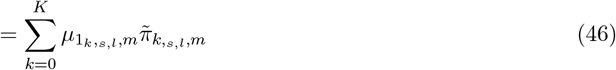

Where 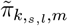is the posterior probability of *κ*_*s,l,m*_

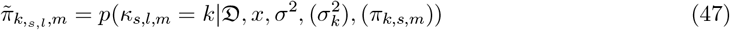

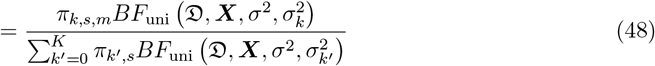

Where *BF*_uni_ is defined as

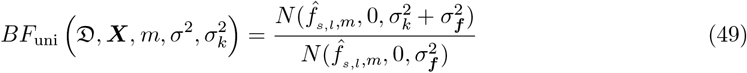

Finally, the Bayes factor for the single multivariate function Bayesian regression using a mixture prior can be computed as follows:

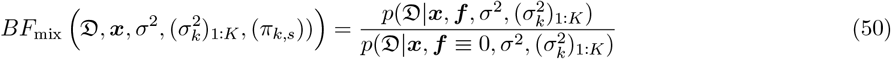

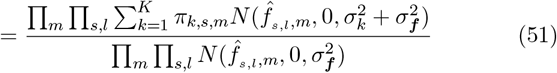

## 3. Bayesian single multiple function

Let’s fit model model 22 in the case where ***X*** is multivariate. Following the seminal work of Wang and colleagues [3] this model can be fitted using the expression described in 33,32, 47, 50 and the posterior distribution of ***γ***_1_

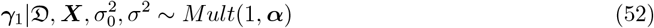

Where the component of the posterior probability of inclusion ***α*** can be computed as following using Bayes rules

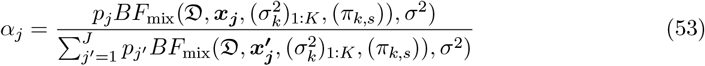

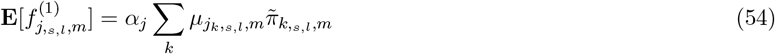

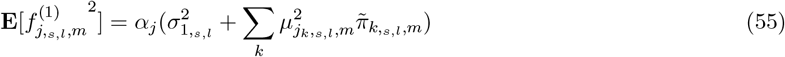

### Estimating hyperparameters (*π*_*k,s,m*_)

As noted earlier, the following *M×K×S*+1 hyperparameters *σ*, (*π*_*k,s,m*_) can be obtained using an empirical Bayes approach. We use a weighted EM algorithm to maximize the marginal likelihood. We provide the detail of the algorithm below. First, notice that the log-likelihood can be written as

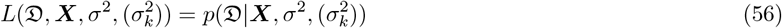

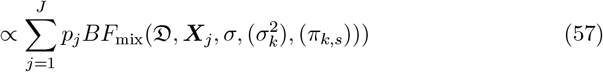

Let *X* be a random variable generated from the following mixture:

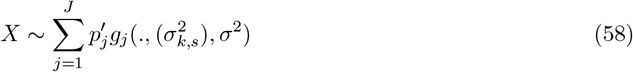

Where 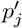 is the proportion of the component *j* in the mixture. Let’s now introduce the latent variable *Z* with value in ⟦1 : *J*⟧, such that 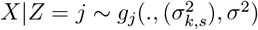and 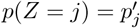

Then the complete log-likelihood can be written

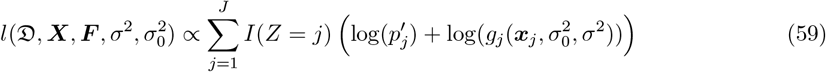

Knowing *Z*|*X*, we can maximize the expected log-likelihood

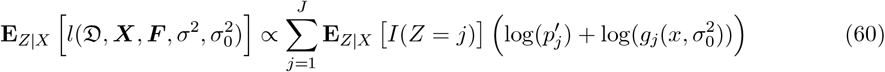

### E step

Thus E step can be computed as

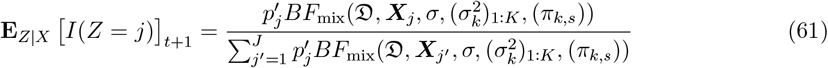

As previously we note ζ_*j,t*+1_ = **E**_*Z*|*X*_ *I*(*Z* = *j*) _*t*+1_

### M step

Thus the complete log-likelihood can be written as

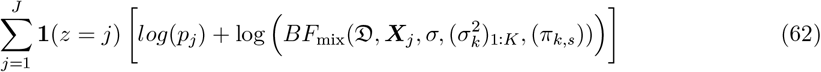

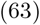

Taking the expected value of the complete log-likelihood is therefore proportional (with respect to (*π*_*k,s*_)) to

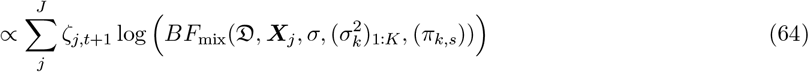

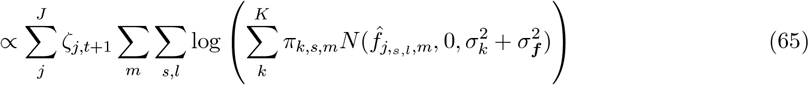

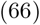

taking the log

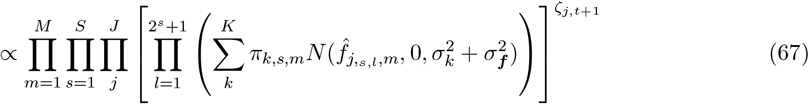

Where 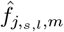 is the maximum likelihood estimate of the j^*th*^ on wavelet coefficient *s, l* in modality *m*. Equation 67 can be maximised using a weighted EM algorithm to optimise over on set of (*π*_*k,s,m*_)_1:*K*_ (here s is fixed) so corresponding *S* × *M* independent weighted maximization problems

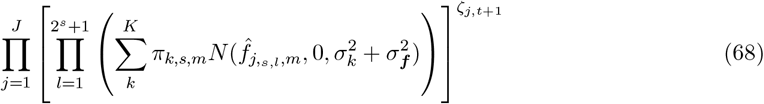

Given that the only unknown quantities are the mixture proportions, these problems are convex optimization problems [39] that can be solved efficiently using interior point methods. In practice we use the *mixsqp R* package to perform these optimization [39].

## 4. Variational algorithm

In this section, we describe our variational approach to fit the sum of single functions model (see equation 18). Let’s rewrite 18 in the following form form:

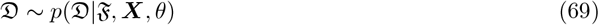

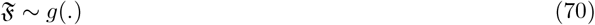

Where D is the matrix of the observed data, ***X*** is the matrix of the observed covariates, F is the latent/unobserved variable of interest, *g* ∈ 𝒢 is a prior distribution for F and *θ* ∈ Θ is the set of the additional/nuisance parameters to be estimated.

We adopt an Empirical Bayes (EB) approach, as we are estimating our prior *g* and the noise parameters *θ* by maximizing the marginal log-likelihood:

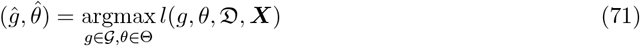

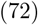

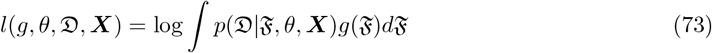

Once these quantities are estimated, we compute the posterior distribution for F :

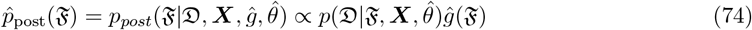

### 4.1. KL divergence and ELBO

As computing the full posterior of model 18 in an intractable problem, we use a VA. More precisely, we seek at finding a distribution *q* that belongs to a certain class of function 𝒬 (detailed in equation 78) that minimizes the KL divergence with the true posterior of model 18. For the sake of pedagogy, let’s rewrite the KL divergence in terms of evidence and evidence lower bound

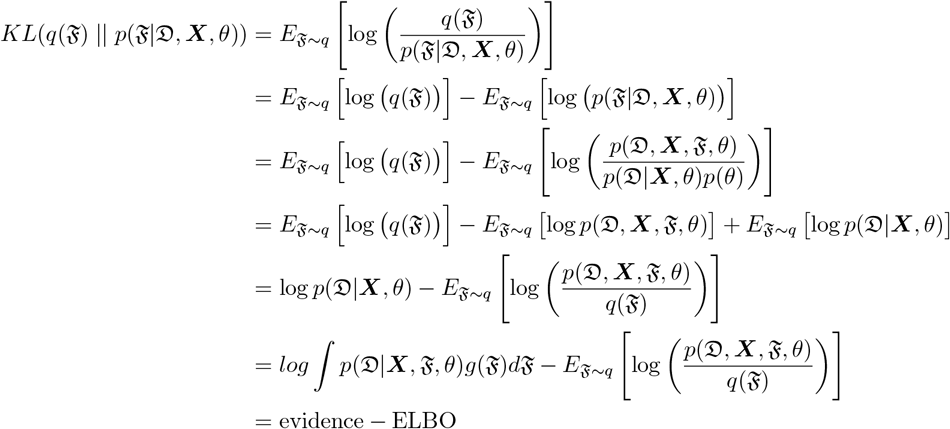

In particular, the ELBO can be written as:

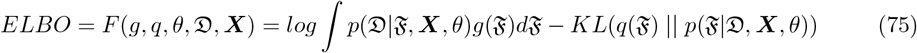

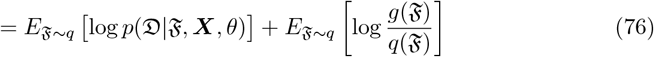

So the ELBO depends on three “parameters” *g*(.), *q*(.), *θ*. Our general optimization problem is the following:

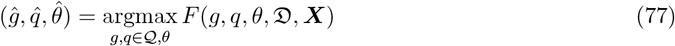

Maximizing the ELBO with respect to *g*(.), *θ* corresponds to our EB procedure (as the first terms of eq 76 corresponds to the marginal likelihood); maximizing the ELBO with respect to *q* ∈ 𝒬 is the “standard” variational Bayes (VB) approach. We optimize 75 in a iterative fashion, by keeping *q*(.) constant and maximize over *g*(.), *θ*. Then by keeping *g*(.), *θ* constant and maximize over *q*(.). The EB procedure to maximize *g* is described in equation 67 . Here we solely focus on maximizing the ELBO while keeping *g* and *θ* constant.

### 4.2. Variational approximation

As previously mentioned, computing the full posterior of 75 is a (likely) intractable problem. Thus, we restrict the form of the “candidate” posterior (*q*) to facilitate the optimization. More precisely, we choose the classical mean-field approximation, so *Q* is defined as

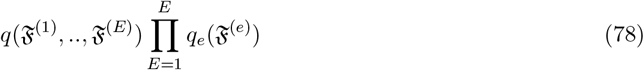

**Form of the ELBO** For *q* ∈ 𝒬 the ELBO can be written as:

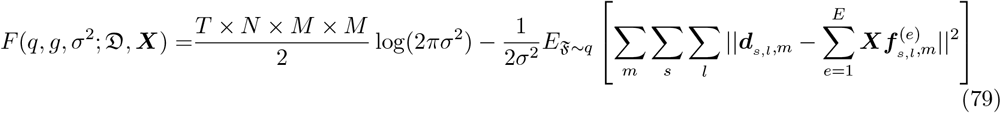

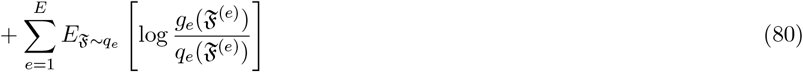

Where || . || is the euclidean norm and ***g*** = (*g*_1_, …, *g*_*E*_) is a collection of mixture prior.

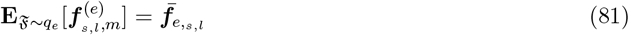

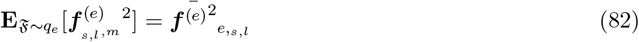

which we provide analytical form in 54 55 given that *g*_*e*_ is a scale mixture prior. Thus the “expected residual sum of squares” (ERSS) can be written as:

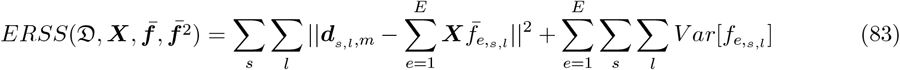

### 4.3 Updating q_e_ and g_e_ in sum of single functions model

In the line of [3] (proposition A.1) we show that maximising *F* (*q, g*, σ^2^; *D*, ***X***) over (*q*_*e*_, *g*_*e*_) corresponds to maximising the ELBO for a simple model where the observed responses are replaced by the partial expected residuals. More precisely:

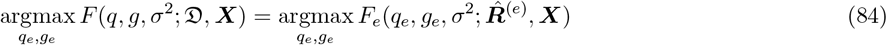

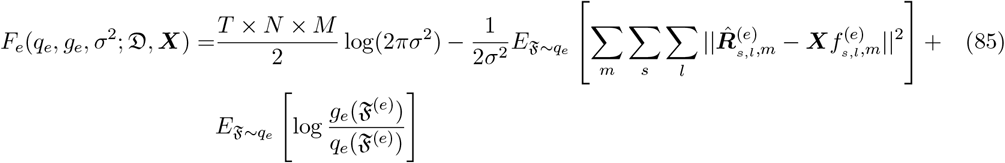

and 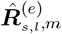 corresponds to the expected residual of the *s, l* wavelet coefficient in frame *m* without effect *e*, i.e.:

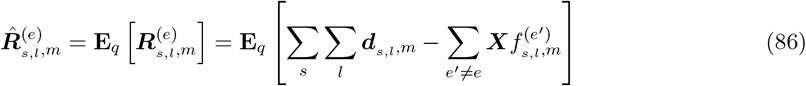

We sometime abuse the notation by noting 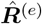 the set of partial residuals wavelet coefficients across all wavelet indexes (*s, l*) and all modalities (*m*).

**Proof** : We rewrite the ELBO only in function of *q*_*e*_, *g*_*e*_, the other terms are captured by *C*, a numerical constant

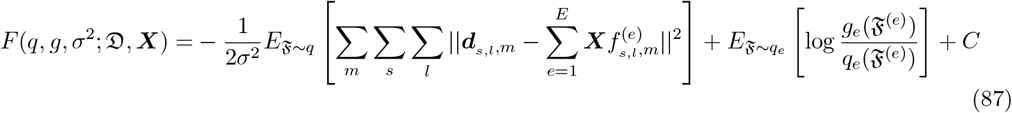

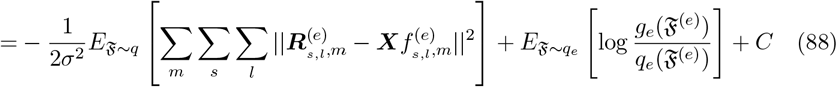

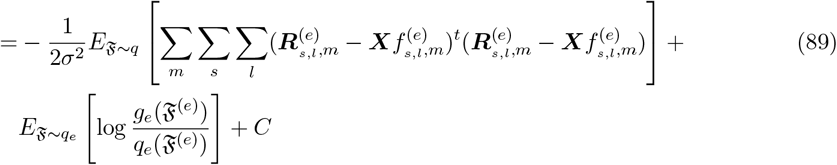

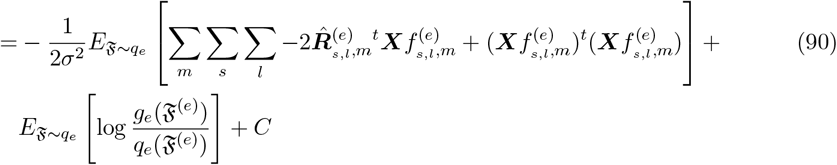

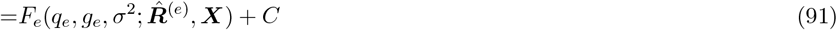

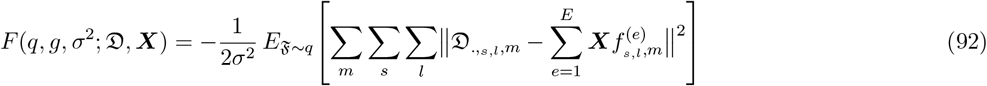

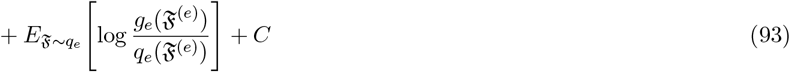

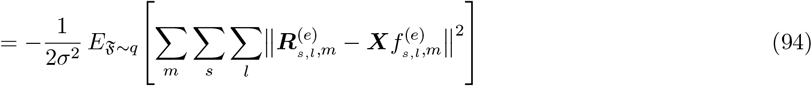

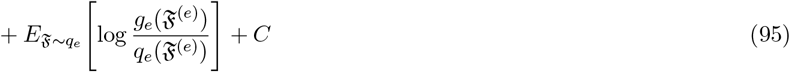

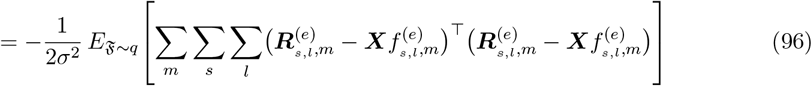

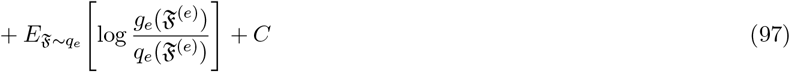

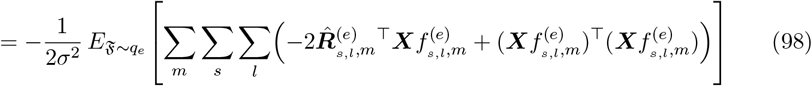

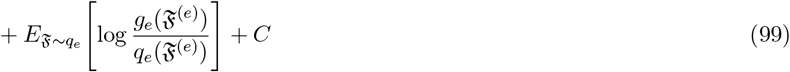

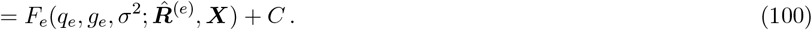

### 4.4. Computing the ELBO

Using equations 54 and 55, the first two components of the ELBO 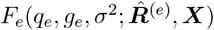 are straightforward to compute (see equation 85). The third term can be computed using the marginal log-likelihood of the single function model for effect *e*; given that 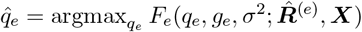, from equation84 it follows that:

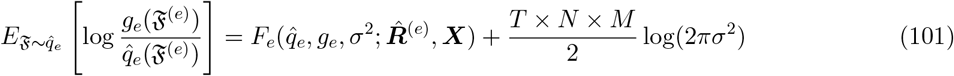

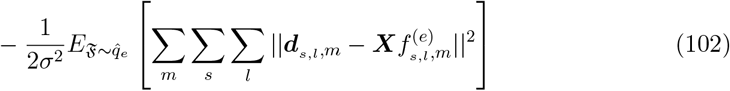

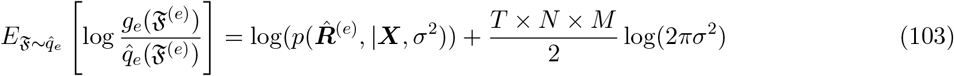

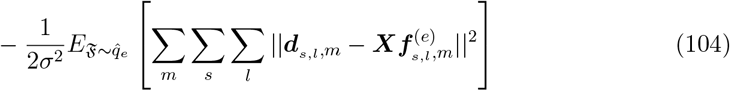

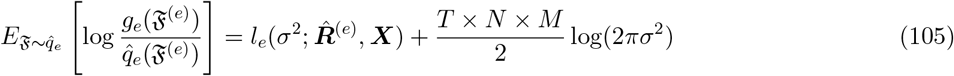

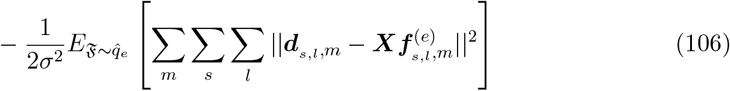

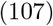

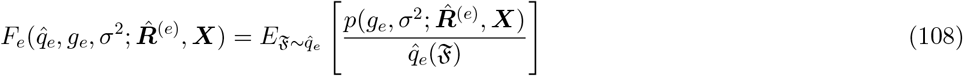

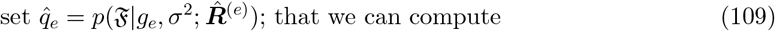

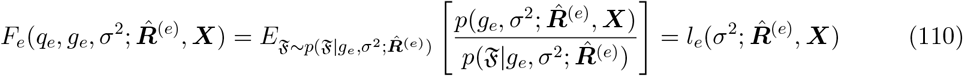

Where 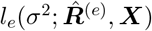 is the log marginal likelihood of the single function model for effect *e*. Furthermore, by definition 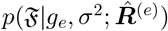 is the overall maximizer over *q*_*e*_ of 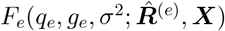. Thus completing the proof.

## References

[1] Urbut, S. M., G. Wang, P. Carbonetto, and M. Stephens (2019, January). Flexible statistical methods for estimating and testing effects in genomic studies with multiple conditions. Nature Genetics 51 (1), 187–195.

[2] Zou, Y., P. Carbonetto, D. Xie, G. Wang, and M. Stephens (2023). Fast and flexible joint fine-mapping of multiple traits via the Sum of Single Effects model. bioRxiv, doi:10.1101/2023.04.14.536893.

[3] Wang, G., A. Sarkar, P. Carbonetto, and M. Stephens (2020). A simple new approach to variable selection in regression, with application to genetic fine mapping. Journal of the Royal Statistical Society, Series B 82 (5), 1273–1300.

[4] Hormozdiari, F., E. Kostem, E. Y. Kang, B. Pasaniuc, and E. Eskin (2014, 8). Identifying causal variants at loci with multiple signals of association. Genetics 198 (2), 497–508.

[5] Arvanitis, M., K. Tayeb, B. J. Strober, and A. Battle (2022). Redefining tissue specificity of genetic regulation of gene expression in the presence of allelic heterogeneity. American Journal of Human Genetics 109 (2), 223–239.

[6] Shim, H., Z. Xing, E. Pantaleo, F. Luca, R. Pique-Regi, and M. Stephens (2024). Multiscale Poisson process approaches for detecting and estimating differences from high-throughput sequencing assays. Annals of Applied Statistics 18 (3), 1773–1788.

[7] Denault, W. R., H. Sun, P. Carbonetto, A. Liu, P. L. De Jager, D. Bennett, G. Wang, M. Stephens, A. D. F. G. Consortium, et al. (2025). fsusie enables fine-mapping of qtls from genome-scale molecular profiles. bioRxiv .

[8] Jaffe, A. E., P. Murakami, H. Lee, J. T. Leek, M. D. Fallin, A. P. Feinberg, and R. A. Irizarry (2012). Bump hunting to identify differentially methylated regions in epigenetic epidemiology studies. International Journal of Epidemiology 41 (1), 200–209.

[9] Pedersen, B. S., D. A. Schwartz, I. V. Yang, and K. J. Kechris (2012). Comb-p: software for combining, analyzing, grouping and correcting spatially correlated P-values. Bioinformatics 28 (22), 2986–2988.

[10] Lent, S., H. Xu, L. Wang, Z. Wang, C. Sarnowski, M.-F. Hivert, and J. Dupuis (2018). Comparison of novel and existing methods for detecting differentially methylated regions. BMC Genetics 19 (Suppl 1), 84.

[11] Peters, T. J., M. J. Buckley, A. L. Statham, R. Pidsley, K. Samaras, R. V Lord, S. J. Clark, and P. L. Molloy (2015). De novo identification of differentially methylated regions in the human genome. Epigenetics & Chromatin 8, 6.

[12] Shim, H. and M. Stephens (2015). Wavelet-based genetic association analysis of functional phenotypes arising from high-throughput sequencing assays. Annals of Applied Statistics 9 (2), 665–686.

[13] Yu, X. and S. Sun (2016). HMM-DM: identifying differentially methylated regions using a hidden Markov model. Statistical Applications in Genetics and Molecular Biology 15 (1), 69–81.

[14] Shen, L., J. Zhu, S.-Y. Robert Li, and X. Fan (2017). Detect differentially methylated regions using non-homogeneous hidden Markov model for methylation array data. Bioinformatics 33 (23), 3701–3708.

[15] Fernàndez, L., M. Pérez, R. Olanda, J. M. Orduña, and J. Marquez-Molins (2020). HPG-DHunter: an ultrafast, friendly tool for DMR detection and visualization. BMC Bioinformatics 21, 287.

[16] Denault, W. R. P. and A. Jugessur (2021). Detecting differentially methylated regions using a fast wavelet-based approach to functional association analysis. BMC Bioinformatics 22, 61.

[17] Lee, W. and J. S. Morris (2016). Identification of differentially methylated loci using wavelet-based functional mixed models. Bioinformatics 32 (5), 664–672.

[18] Schor, I. E., J. F. Degner, D. Harnett, E. Cannavó, F. P. Casale, H. Shim, D. A. Garfield, E. Birney, M. Stephens, O. Stegle, and E. E. M. Furlong (2017). Promoter shape varies across populations and affects promoter evolution and expression noise. Nature Genetics 49 (4), 550–558.

[19] Fusi, N. and J. Listgarten (2017). Flexible modeling of genetic effects on function-valued traits. Journal of Computational Biology 24 (6), 524–535.

[20] Collado-Torres, L., A. Nellore, A. C. Frazee, C. Wilks, M. I. Love, B. Langmead, R. A. Irizarry, J. T. Leek, and A. E. Jaffe (2016). Flexible expressed region analysis for RNA-seq with derfinder. Nucleic Acids Research 45 (2), e9.

[21] Collado-Torres, L., A. Nellore, K. Kammers, S. E. Ellis, M. A. Taub, K. D. Hansen, A. E. Jaffe, B. Langmead, and J. T. Leek (2017). Reproducible RNA-seq analysis using recount2. Nature Biotechnology 35 (4), 319–321.

[22] Zou, Y., P. Carbonetto, G. Wang, and M. Stephens (2022). Fine-mapping from summary data with the “Sum of Single Effects” model. PLoS Genetics 18 (7), e1010299.

[23] Bennett, D. A., A. S. Buchman, P. A. Boyle, L. L. Barnes, R. S. Wilson, and J. A. Schneider (2018). Religious orders study and rush memory and aging project. Journal of Alzheimer’s disease 64 (1), S161–S189.

[24] Denault, W. R. P., H. Sun, P. Carbonetto, A. Liu, P. L. D. Jager, D. Bennett, T. A. D. F. G. Consortium, G. Wang, and M. Stephens (2025, August). fSuSiE enables fine-mapping of QTLs from genome-scale molecular profiles. ISSN: 2692-8205 Pages: 2025.08.17.670732 Section: New Results.

[25] Yuan, K., R. J. Longchamps, A. F. Pardiñas, M. Yu, T.-T. Chen, S.-C. Lin, Y. Chen, M. Lam, R. Liu, Y. Xia, Z. Guo, W. Shi, C. Shen, T. S. W. o. P. G. Consortium, M. J. Daly, B. Neale, Y.-C. A. Feng, Y.-F. Lin, C.-Y. Chen, M. O’Donovan, T. Ge, and H. Huang (2023, January). Finemapping across diverse ancestries drives the discovery of putative causal variants underlying human complex traits and diseases. ISSN: 2328-4293 Pages: 2023.01.07.23284293.

[26] Mallat, S. (1989). A theory for multiresolution signal decomposition: the wavelet representation. IEEE Transactions on Pattern Analysis and Machine Intelligence 11 (7), 674–693.

[27] Mallat, S. G. (2009). A wavelet tour of signal processing: the sparse way (3rd ed.). Boston, MA: Elsevier/Academic Press.

[28] Dimitromanolakis, A., J. Xu, A. Krol, and L. Briollais (2019). sim1000G: a user-friendly genetic variant simulator in R for unrelated individuals and family-based designs. BMC Bioinformatics 20, 26. Fine-mapping molecular QTLs using mfSuSiE 15

[29] Xiong, X., B. T. James, C. A. Boix, Y. P. Park, K. Galani, M. B. Victor, N. Sun, L. Hou, L.-L. Ho, J. Mantero, A. N. Scannail, V. Dileep, W. Dong, H. Mathys, D. A. Bennett, L.-H. Tsai, and M. Kellis (2023). Epigenomic dissection of Alzheimer’s disease pinpoints causal variants and reveals epigenome erosion. Cell 186 (20), 4422–4437.e21.

[30] Leung, Y. Y., W.-P. Lee, A. B. Kuzma, H. Nicaretta, O. Valladares, P. Gangadharan, L. Qu, Y. Zhao, Y. Ren, P.-L. Cheng, et al. (2025). Alzheimer’s disease sequencing project release 4 whole genome sequencing dataset. Alzheimer’s & Dementia 21 (5), e70237.

[31] Wallace, C. (2021). A more accurate method for colocalisation analysis allowing for multiple causal variants. PLoS genetics 17 (9), e1009440.

[32] Zou, Y., P. Carbonetto, D. Xie, G. Wang, and M. Stephens (2024). Fast and flexible joint fine-mapping of multiple traits via the sum of single effects model. bioRxiv, 2023–04.

[33] Bellenguez, C., F. Küçükali, I. E. Jansen, L. Kleineidam, S. Moreno-Grau, N. Amin, A. C. Naj, R. Campos-Martin, B. Grenier-Boley, V. Andrade, et al. (2022). New insights into the genetic etiology of alzheimer’s disease and related dementias. Nature genetics 54 (4), 412–436.

[34] The 1000 Genomes Project Consortium (2015). A global reference for human genetic variation. Nature 526 (7571), 68–74.

[35] Maller, J. B., G. McVean, J. Byrnes, D. Vukcevic, K. Palin, Z. Su, et al. (2012). Bayesian refinement of association signals for 14 loci in 3 common diseases. Nature Genetics 44 (12), 1294– 1301.

[36] Sniekers, S. and A. van der Vaart (2020). Adaptive Bayesian credible bands in regression with a Gaussian process prior. Sankhya A 82 (2), 386–425.

[37] Barber, S., G. P. Nason, and B. W. Silverman (2002). Posterior probability intervals for wavelet thresholding. Journal of the Royal Statistical Society, Series B 64 (2), 189–205.

[38] Wakefield, J. (2009). Bayes factors for genome-wide association studies: comparison with P-values. Genetic Epidemiology 33 (1), 79–86. eprint: https://onlinelibrary.wiley.com/doi/pdf/10.1002/gepi.20359.

[39] Kim, Y., P. Carbonetto, M. Stephens, and M. Anitescu (2020). A fast algorithm for maximum likelihood estimation of mixture proportions using sequential quadratic programming. Journal of Computational and Graphical Statistics 29 (2), 261–273.

## Methods-only references

[40] Fachal, L., H. Aschard, J. Beesley, D. R. Barnes, J. Allen, S. Kar, et al. (2020). Fine-mapping of 150 breast cancer risk regions identifies 191 likely target genes. Nature Genetics 52, 56–73.

[41] Yang, Z., C. Wang, L. Liu, A. Khan, A. Lee, B. Vardarajan, R. Mayeux, K. Kiryluk, and I. Ionita-Laza (2023). CARMA is a new Bayesian model for fine-mapping in genome-wide association meta-analyses. Nature Genetics 55 (6), 1057–1065.

[42] Ma, L. and J. Soriano (2018). Efficient functional ANOVA through wavelet-domain Markov Groves. Journal of the American Statistical Association 113 (522), 802–818.

[43] Crouse, M., R. Nowak, and R. Baraniuk (1998). Wavelet-based statistical signal processing using hidden Markov models. IEEE Transactions on Signal Processing 46 (4), 886–902.

[44] Nason, G. (2008). Wavelet Methods in Statistics with R. New York, NY: Springer.

